# A systems genetics approach reveals environment-dependent associations between SNPs, protein co-expression and drought-related traits in maize

**DOI:** 10.1101/636514

**Authors:** Mélisande Blein-Nicolas, Sandra Sylvia Negro, Thierry Balliau, Claude Welcker, Llorenç Cabrera Bosquet, Stéphane Dimitri Nicolas, Alain Charcosset, Michel Zivy

## Abstract

The effect of drought on maize yield is of particular concern in the context of climate change and human population growth. However, the complexity of drought-response mechanisms make the design of new drought-tolerant varieties a difficult task that would greatly benefit from a better understanding of the genotype-phenotype relationship. To provide novel insight into this relationship, we applied a systems genetics approach integrating high-throughput phenotypic, proteomic and genomic data acquired from 254 maize hybrids grown under two watering conditions. Using association genetics and protein co-expression analysis, we detected more than 22,000 pQTLs across the two conditions and confidently identified fifteen loci with potential pleiotropic effects on the proteome. We showed that even mild water deficit induced a profound remodeling of the proteome, which affected the structure of the protein co-expression network, and a reprogramming of the genetic control of the abundance of many proteins, notably those involved in stress response. Co-localizations between pQTLs and QTLs for ecophysiological traits, found mostly in the water deficit condition, indicated that this reprogramming may also affect the phenotypic level. Finally, we identified several candidate genes that are potentially responsible for both the co-expression of stress response proteins and the variations of ecophysiological traits under water deficit. Taken together, our findings provide novel insights into the molecular mechanisms of drought tolerance and suggest some pathways for further research and breeding.

## INTRODUCTION

Maize is the world’s leading crop (Shiferaw et al. 2011) in terms of production. Having a C4 metabolism, it exhibits high water use efficiency (WUE). However, it is also highly susceptible to water deficit. For example, maize is more affected by drought than either its close relative sorghum (above-ground dry biomass reduced by 47-51 % and 37-38 %, respectively; Zegada-Lizarazu et al. 2012, Schittenhelm and Schroetter 2014) or wheat, which is a C3 plant (yield reduction associated with a 40% water reduction of 40% and 20.6%, respectively; Daryanto et al. 2016). Improving maize yield under drought has been an important aim of breeding programs for decades (Campos et al. 2004, 2006; Cooper et al. 2014). However, despite the overall genetic improvement of maize, increases in drought sensitivity have been reported in several regions (Lobell et al. 2014; Zipper et al. 2016; Meng et al. 2016). In addition, severe episodes of drought are projected to become more frequent in the near future due to climate change (Harrison et al. 2014). Therefore, maize productivity under water deficit is of particular concern and large efforts are still required to design varieties that are able to maintain high yields in drought conditions.

One lever to accelerate crop improvement is to better understand the genetic and molecular bases of drought tolerance. This highly complex trait is associated with a series of mechanisms occurring at different spatial and temporal scales to (i) stabilize the plant’s water and carbon status, (ii) control the side effects of water deficit including oxidative stress, mineral deficiencies and reduced photosynthesis and (iii) maintain plant yield (Chaves et al. 2003). At the physiological level, short-term responses include stomata closure, adjustment of osmotic and hydraulic conductance, leaf growth inhibition and root growth promotion (Tardieu et al. 2018). At the molecular level, complex signaling and regulatory events occur, involving several hormones, of which abscisic acid (ABA) is a key player, and a broad range of transcription factors (Golldack et al. 2014; Osakabe et al. 2014; Tripathi et al. 2014). Molecular responses also include the accumulation of metabolites involved in osmotic adjustment, membrane and protein protection, as well as scavenging of reactive oxygen species, and the expression of drought-responsive proteins such as dehydrins, late embryogenesis abundant (LEA) and heat shock proteins (HSP) (Valliyodan and Nguyen 2006; Seki et al. 2007). All these responses will depend on the drought scenario, the phenological stage, the genetic makeup and the general environmental conditions (Tardieu et al. 2018). Taken together, the multiplicity and versatility of the mechanisms involved explain the difficulty in selecting for drought tolerance.

A better understanding of the genotype-phenotype relationship will help guide the development of new drought-tolerant varieties. Systems genetics is a recent approach providing improved insight into this relationship by deciphering the biological networks and molecular pathways underlying complex traits and by investigating how these traits are regulated at the genetic and epigenetic levels (Nadeau and Dudley 2011; Civelek and Lusis 2014; Feltus 2014; van der Sijde et al. 2014; Markowetz and Boutros 2015). This approach compares the position of quantitative trait loci (QTLs) underlying phenotypic traits variation to that of QTLs underlying the variation of upstream molecular traits such as transcript expression (eQTLs) or protein abundance (pQTLs). Until recently, this approach had been mostly applied in human and mice (Moreno-Moral and Petretto 2016) In plants, systems genetics studies have been carried out in a few species including wheat (Munkvold et al. 2013), rapeseed (Basnet et al. 2016), eucalyptus (Mizrachi et al. 2017) and maize (Christie et al. 2017; Jiang et al. 2019).

The first studies comparing QTLs and pQTLs used 2D gel proteomics to quantify proteins (Bourgeois et al. 2011; de Vienne et al. 1999). Since then, proteome coverage and data reliability have been widely improved by the use of mass spectrometry (MS)-based proteomics (Wasinger et al. 2013). Despite these technical advancements, the systems genetics studies published so far have preferentially used transcripts rather than proteins as the intermediate level between the genome and phenotypic traits. One reason is that large-scale proteomics experiments remain challenging (Blein-Nicolas et al. 2015) due to technical constraints (Balliau et al. 2018) and the trade-off between depth of coverage and sample throughput (Keshishian et al. 2017). However, proteins are particularly relevant molecular components for linking genotype to phenotype. Indeed, proteins abundance is expected to be more highly related to phenotype than transcript expression due to the buffering of transcriptional variations and the role of post-translational regulations in phenotype construction (Foss et al. 2011; Battle et al. 2015; Chick et al. 2016; Albertin et al. 2013; Vogel and Marcotte 2012).

Here, we aimed to better understand the molecular mechanisms associated with the genetic polymorphisms underlying the variations of ecophysiological traits related to drought tolerance. To this end, we performed a novel systems genetics study where MS-based proteomics data acquired from 254 maize genotypes grown in two watering conditions were integrated with high-throughput genomic and phenotypic data. First, protein abundance was analyzed using a genome wide association study (GWAS) and co-expression networks. Then, these data were integrated with ecophysiological phenotypic data from the same experiment (Prado et al. 2018) using a correlation analysis and by searching for QTL/pQTL co-localizations.

## RESULTS

### Mild water deficit has extensively remodeled the proteome

Using MS-based proteomics, we analyzed more than 1,000 leaf samples taken from 254 genotypes representing the genetic diversity within dent maize and grown in well-watered (WW) and water deficit (WD) conditions. After data filtering, the peptide intensity dataset included 977 samples corresponding to 251 genotypes from which we quantified 2,055 proteins described in Supplemental Table S1. Among these, 973 were quantified by integration of peptide intensities (XIC-based quantification). The remaining 1,082, whose peptides had more than 10% of missing intensity values, were quantified by spectral counting (SC-based quantification). Note that the latter proteins were less abundant (Supplemental Fig. S1A) and less precisely quantified (Supplemental Fig. S1B) than those that could be quantified by XIC.

Heatmap representations of protein abundance showed that two large separate protein clusters were associated with the two watering conditions (Fig. 1A). This indicates that, although moderate, water deficit had extensively remodeled the proteome of most genotypes. Accordingly, 82.4% and 74.5% of proteins from the XIC-based and SC-based sets, respectively, responded significantly to water deficit (Supplemental Table S1, Supplemental Figure S2A). These included several proteins known to be involved in responses to drought or stress (Shinozaki & Yamaguchi-Shinozaki 2007; Wang et al. 2016) such as dehydrins (GRMZM2G079440, GRMZM2G373522), an ABA-responsive protein (GRMZM2G106622), an LEA protein (GRMZM2G352415), HSPs (GRMZM2G360681, GRMZM2G080724, GRMZM2G112165), a phospholipase D (GRMZM2G061969), a glyoxalase I (GRMZM2G181192) and a gluthathione-S-transferase (GRMZM2G043291). Induced and repressed proteins constituted two highly differentiated populations in terms of function (Fig. 1B). In particular, transcription, translation, energy metabolism and metabolism of cofactors and vitamins were better represented within repressed proteins, while carbohydrate and amino acid metabolism and environmental adaptation were better represented within induced proteins.

**Figure 1.**
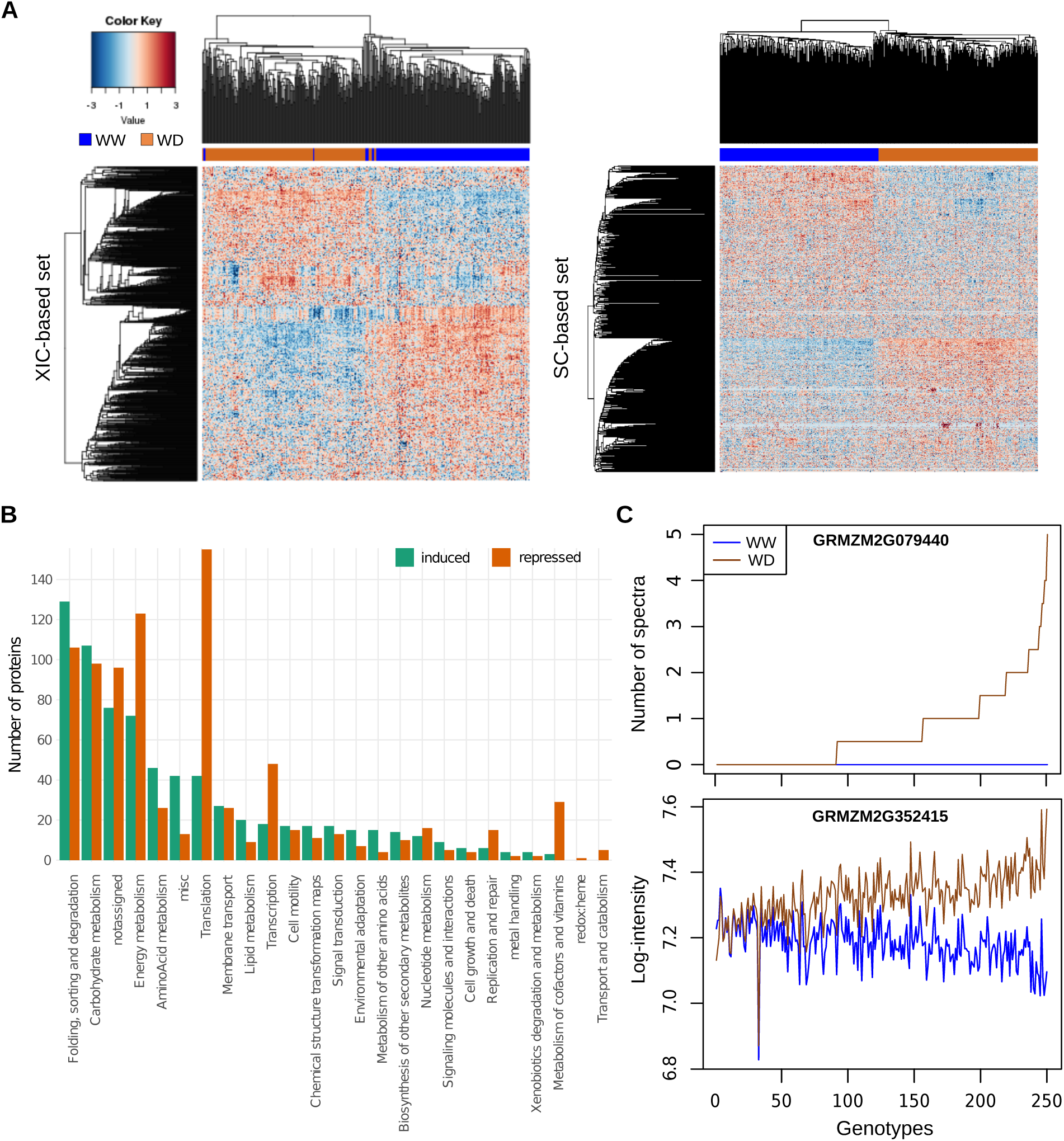
Effect of a mild water deficit on the proteome. (A) Heatmap representations of protein abundances estimated for the XIC-based protein set (left) and the SC-based protein set (right). Each line corresponds to a protein and each column to a genotype x watering condition combination. For each protein, abundance values were scaled and represented by a color code as indicated by the color-key bar. Hierarchical clustering of the genotype x watering condition combinations (top) and of the proteins (left) was built using the 1-Pearson correlation coefficient as the distance and the unweighted pair group method with arithmetic mean (UPGMA) as the aggregation method. (B) Functions of induced and repressed proteins under water deficit. (C) Abundance profiles of the RAB17 dehydrin (GRMZM2G079440 quantified based on the number of spectra) and of a LEA protein (GRMZM2G352415 quantified based on peptide intensity) in the two watering conditions. Genotypes on the x axis were ordered according to the WD/WW abundance ratio. The lists of genotypes in this order are available in Supplemental Table S2.

The global impact of genotypic change on the proteome was less extensive than that of water deficit, since the proteomes of two different genotypes grown in the same watering condition were more similar than the proteomes of a same genotype grown in different conditions (Fig. 1A). However, the maximum amplitudes of abundance variations were similar (Supplemental Fig. S2B). In addition, 94.9% of proteins from the XIC-based set exhibited significant variation in abundance attributable to genetic variation (Supplemental Table S1). This was confirmed by broad sense heritability, the median of which was 0.47 and 0.46 for WW and WD conditions, respectively (Supplemental Fig. S3A). By contrast, in the SC-based set, only 34.4% of the proteins showed significant variation in abundance attributable to genetic variation with a median broad sense heritability of 0.08 and 0.10 for WW and WD conditions, respectively (Supplemental Fig. S3B).

Significant GxE interactions were detected for only four and 12 proteins from the SC-based and XIC-based set, respectively, probably due to a lack of statistical power. These proteins included an LEA protein (GRMZM2G352415), a HSP (GRMZM2G083810_P01) and a COR410 dehydrin (GRMZM2G147014_P01). Although the GxE interaction was not statistically significant for the dehydrin ZmRab17 (GRMZM2G079440), this protein was remarkable because it was undetectable in the WW condition and more or less expressed, depending on genotype, in the WD condition (Fig. 1C, Supplemental Table S2).

### The strength of the genetic control of protein abundance is related to protein function

We performed GWAS for 2,501 combinations of protein abundance x watering condition showing a heritability > 0.2. In total, we detected 514,270 significant associations for 2,466 (98.6%) combinations of protein abundance x watering condition involving 1,367 proteins. When summarizing associated SNPs into pQTLs using classical methods based on genetic distance or linkage disequilibrium (LD), we observed a positive relationship between the number of pQTLs per chromosome and the *P-value* of the most strongly associated pQTL from the corresponding chromosome (Supplemental Fig. S4A,B). To get rid of this artefactual relationship, which could lead to the detection of more than 250 pQTLs on one chromosome, we developed a geometric method based on the *P-value* signal of SNPs (Supplemental Fig. S4C). This method produced the lowest number of pQTLs per combination of protein abundance x watering condition (median = 8 *vs* 13 for the two other methods) and the lowest maximum number of pQTLs per chromosome (18 *vs* 272 and 209 for the methods based on genetic distance and LD, respectively). Using this geometric method and considering only pQTLs accounting for more than 3% of the total variance, we thus detected 22,664 pQTLs accounting for 3 to 77.1% of the variance (Supplemental Table S3). Of these, 1,113 were local, *i.e.* located less than 10^6^ bp from the protein encoding gene, of which 339 were located within the genes. Among distant pQTLs, 80.9% were located on a different chromosome from that of the protein encoding gene. Local pQTLs had stronger effects than distant pQTLs (average *R2*=15.3% and 5.2%, respectively; Supplemental Fig. S5). For 485 proteins, no local pQTL was detected in either condition. This set of proteins was significantly enriched in proteins involved in translation (15.3% *vs* 3.8%, adjusted *P-value* = 2.3E^-10^) and energy metabolism (17.3% *vs* 8.5%, adjusted *P-value* = 8.2E^-5^) and depleted in proteins involved in carbohydrate metabolism (9.9% *vs* 19.5%, adjusted *P-value* = 8.2E^-5^) compared to the 662 proteins showing a local pQTL in at least one condition. They also exhibited fewer distant pQTLs and were much less heritable (Supplemental Fig. S6). These results indicate that the strength of the genetic control over protein abundance depends on protein function. This observation is supported by the positive correlation between the mean number of pQTLs and the mean heritability per functional category (Fig. 2).

**Figure 2.**
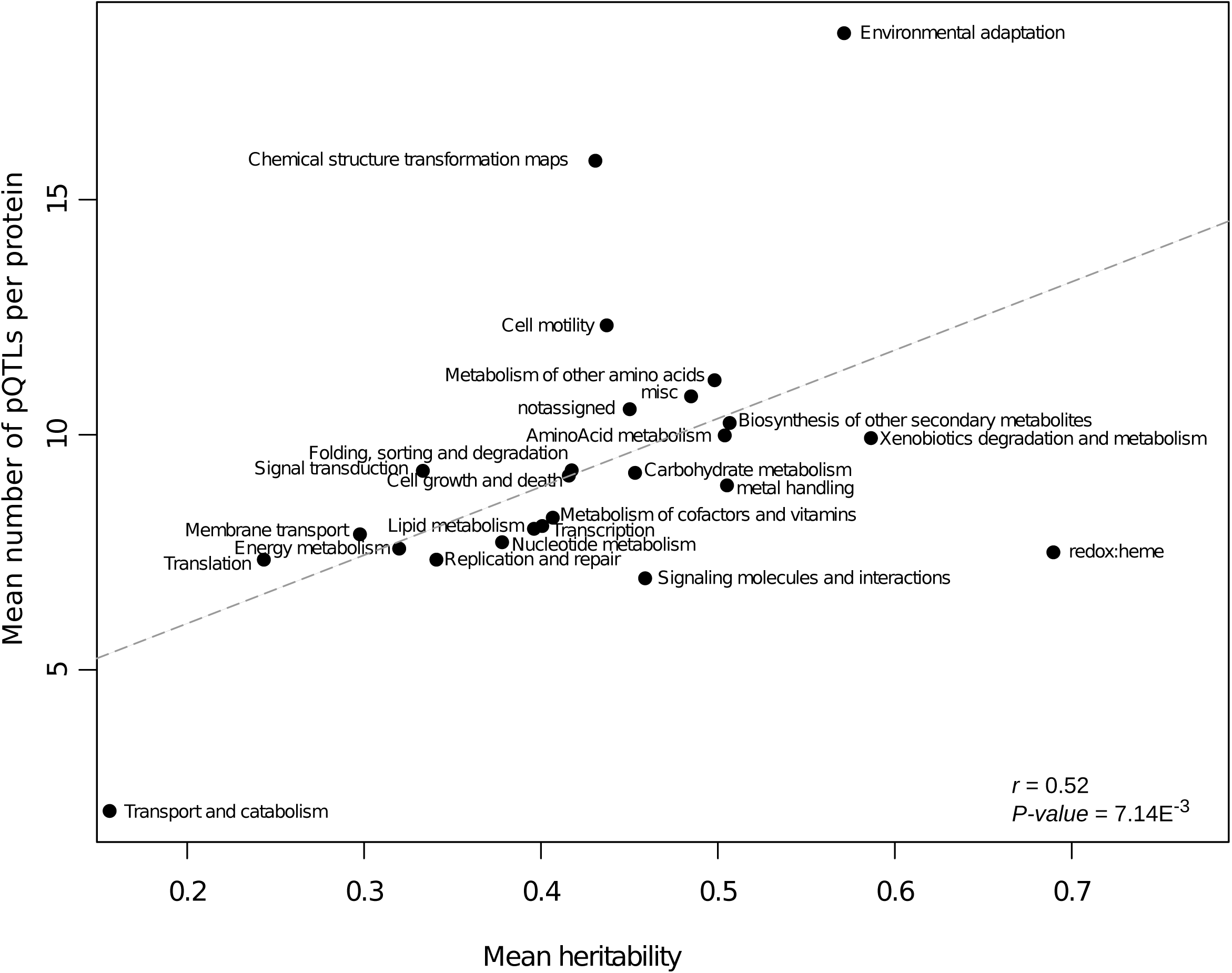
Relationship between the mean number of pQTLs per KEGG category and the mean heritability per KEGG category.

### Identification of loci with potential pleiotropic effects on the proteome

pQTLs were not uniformly distributed in the genome (Fig. 3). Instead, there were genomic regions enriched with pQTLs. We detected 26 and 31 such hotspots that contained at least 19 pQTLs in the WW and WD conditions, respectively (Supplemental Table S4). These hotspots may represent loci with pleiotropic effects on the proteome, *i.e.* loci associated with the abundance variation of several proteins. To refine the detection of such loci, we used a second independent approach based on the search for co-expression QTLs (coQTLs), *i.e.*, QTLs associated to the abundance variations of several co-expressed proteins. To do so, we first performed a weighted gene co-expression network analysis (WGCNA) of protein co-expression across the 251 genotypes in the two watering conditions separately (Supplemental Table S5). The two resulting networks differed in the presence of condition-specific modules indicating that water deficit has altered the structure of the protein co-expression network (Fig. 4, Supplemental Fig. S7, Supplemental File S1). For each co-expression module, we then submitted the representative variable, called eigengene according to the WGCNA terminology, to GWAS in order to identify coQTLs. In total, we detected 176 coQTLs (96 for the 8 WW modules and 80 for the 8 WD modules, Supplemental Table S3). Fifteen of them co-localized with pQTL hotspots (Supplemental Table S4). Thus by crossing these results, we confidently identified four loci in the WW condition and eleven loci in the WD condition as having potential pleiotropic effects on the proteome. Of note, the proteins that were associated with hotspots Hs22d and Hs21d and that were also in the modules having coQTLs co-localising with these hotspots were mainly ribosomal proteins (Supplemental Table S6). This suggests that Hs22d and Hs21d may contain loci involved in ribosome biogenesis.

**Figure 3.**
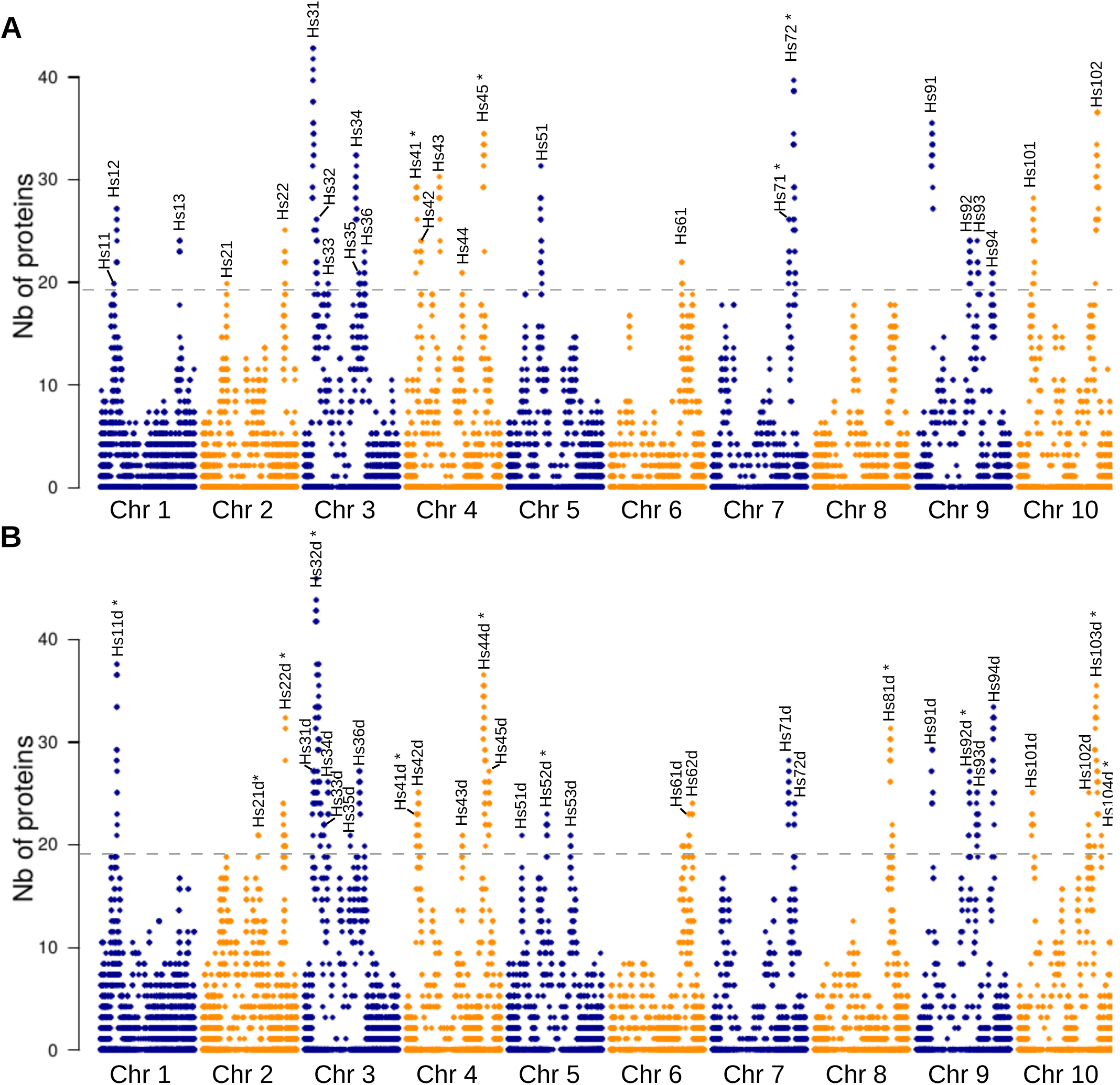
Distribution of pQTLs across the genome. (A) In the well-watered condition. (B) In the water deficit condition. Each point indicates the number of proteins controlled by a pQTL located within a given genomic region defined by the linkage disequilibrium interval around an SNP. Dashed horizontal lines indicate the threshold used to detect pQTL hotspots. Names and positions of the pQTL hotspots are indicated above each graph. Asterisks indicate the pQTL hotspots confidently detected as loci with potential pleiotropic effects (details given in Supplemental Table S4).

**Figure 4.**
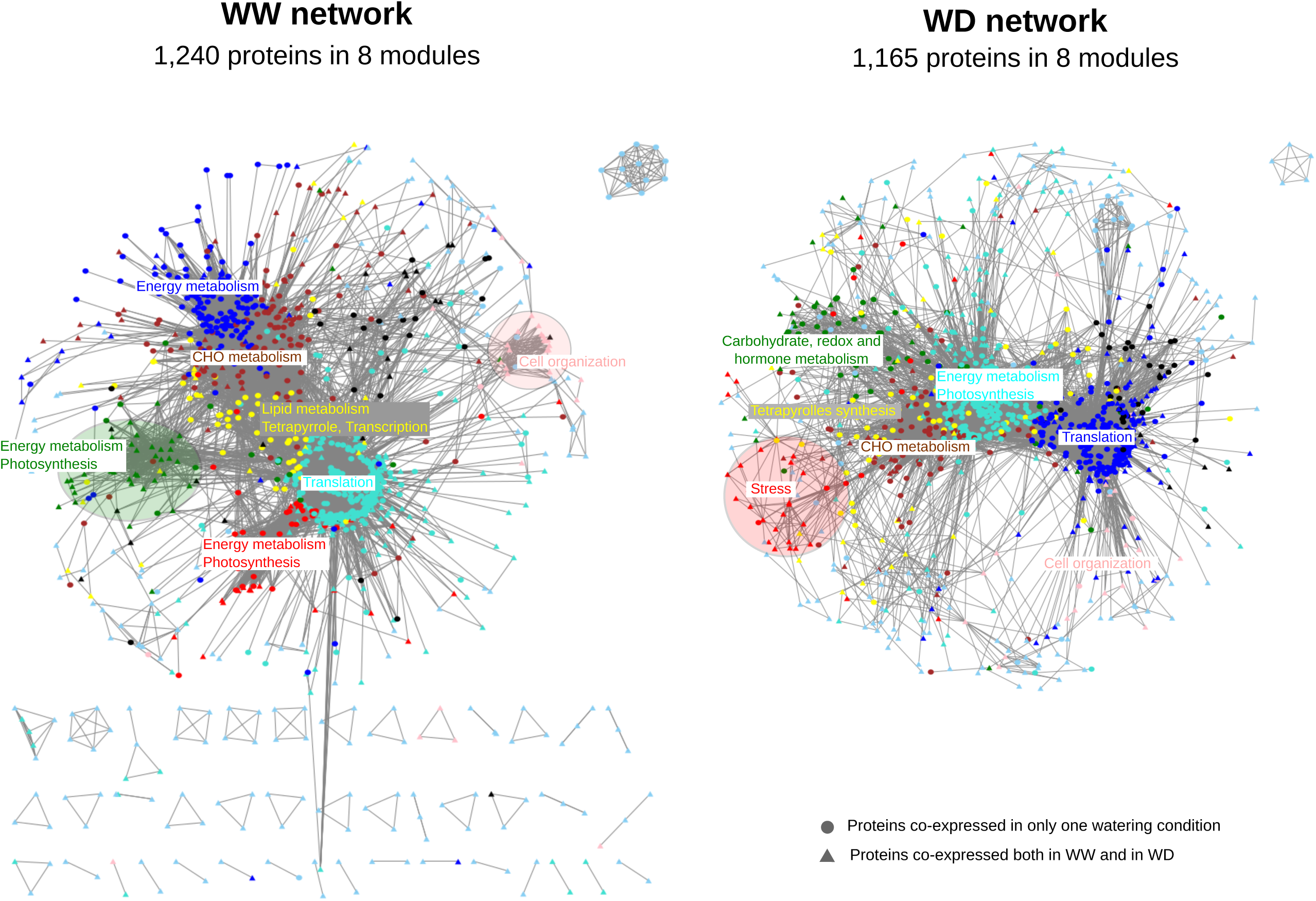
Graphical representation of the co-expression networks resulting from the WGCNA analysis. Only proteins with an adjacency > 0.02 are shown. The two views were created by Cytoscape v3.5.1 using an unweighted, spring-embedded layout (cytoscape files are available in Supplementary File S1). The colors displayed on each network represent the different modules identified by WGCNA. Functional enrichments of modules are indicated with corresponding colors. Condition-specific modules are circled. Each module contains 35 to 471 proteins.

### The genetic architecture of protein abundances depends on the environment

Of the 11,034 pQTLs detected in the WW condition, only 1,124 (10.2%) had a co-localizing pQTL in the WD condition. These pQTLs were generally of strong effect (Supplemental Fig. S8A) and were enriched in local pQTLs (32.6% *vs* 4.9% over the entire dataset). While most of the pQTLs that were common to the two conditions had similar effects in both conditions, 75 (6.7%) of them exhibited contrasted effects (Supplemental Fig. S8B). Half of these pQTLs were local, suggesting that gene promoters may be involved in the GxE interaction or that the pQTLs that were detected in each condition corresponded to different polymorphic sequences with different effects on protein abundance. These pQTLs were associated with 70 proteins, several of which were stress-responsive (*e.g.* the LEA protein GRMZM2G045664, HSPs GRMZM5G813217 and GRMZM2G536644, Supplemental Table S7). Altogether, these results show that water deficit has altered the genetic architecture of protein abundance.

### Identification of loci associated with trait variation at multiple scales

To gain insight into the molecular mechanisms associated with drought tolerance, we searched for co-localizations between the pQTLs, coQTLs and hotspots detected in our study and the 160 QTLs identified by Prado et al. (2018) on the same plant material. These QTLs were associated with eight ecophysiological traits related to growth and transpiration rate: early leaf area (*i.e.* before water deficit; LAe), late leaf area (LAl), early biomass (Be), late biomass (Bl), water use (WU), water use efficiency (WUE), stomatal conductance (gs) and transpiration rate (Trate). Robust co-localizations were determined by taking into account the correlation between each trait and protein values.

In total, we identified 68 pairs of SNPs corresponding to QTL/pQTL co-localizations (Fig. 5, Supplemental Table S8). Only one involved a local pQTL. The QTL/pQTL distance was generally less than 100 kb, with, in 25.7% of cases, the same SNP representing the QTL and the pQTL (Fig. 6A). Most QTL/pQTL co-localizations (98%) were detected in the WD condition, where they corresponded to 39 of the 91 QTLs reported in this condition (Prado et al. 2018). They involved six ecophysiological phenotypic traits (Bl, LAl, WU, WUE, Trate and gs) and 47 proteins, many of which were stress-responsive (Supplemental Table S9). Twenty-three proteins exhibited multiple QTL/pQTL co-localizations (Supplemental Table S9).

**Figure 5.**
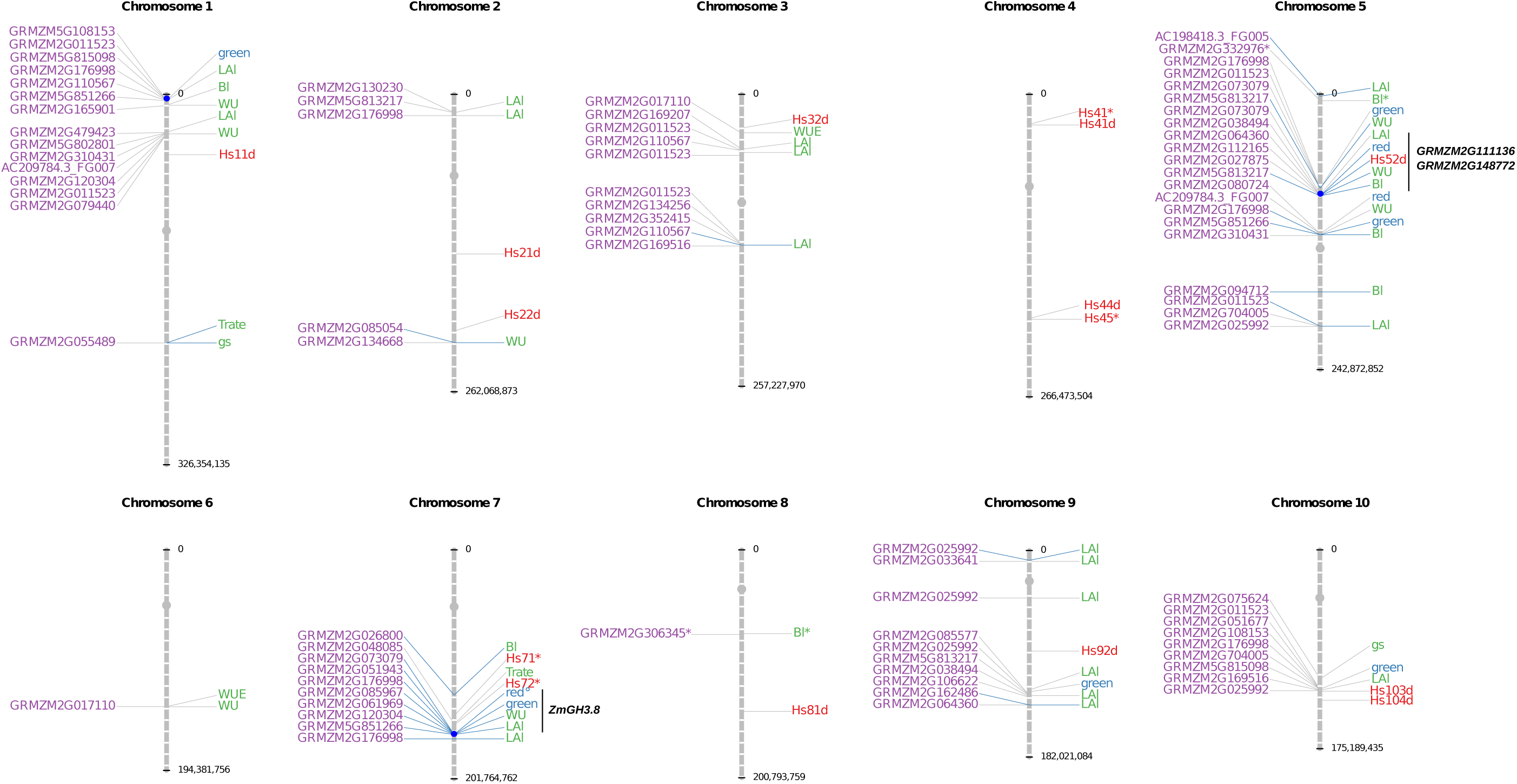
Genomic positions of the co-localizing pQTLs, coQTLs and QTLs. The positions of the fifteen pQTL hotspots confidently identified as loci with potential pleiotropic effects are indicated, as well as the position of the most promising candidate genes. Chromosomes are segmented into 10 Mb bins. Grey dots represent the centromeres and blue dots indicate the position of genomic regions showing evidences for pleiotropy at both the proteome and phenotype level. Blue lines indicate co-localizations with QTLs that are determined by a same SNP. ° WD-specific module, * co-localization found in the WW condition.

**Figure 6.**
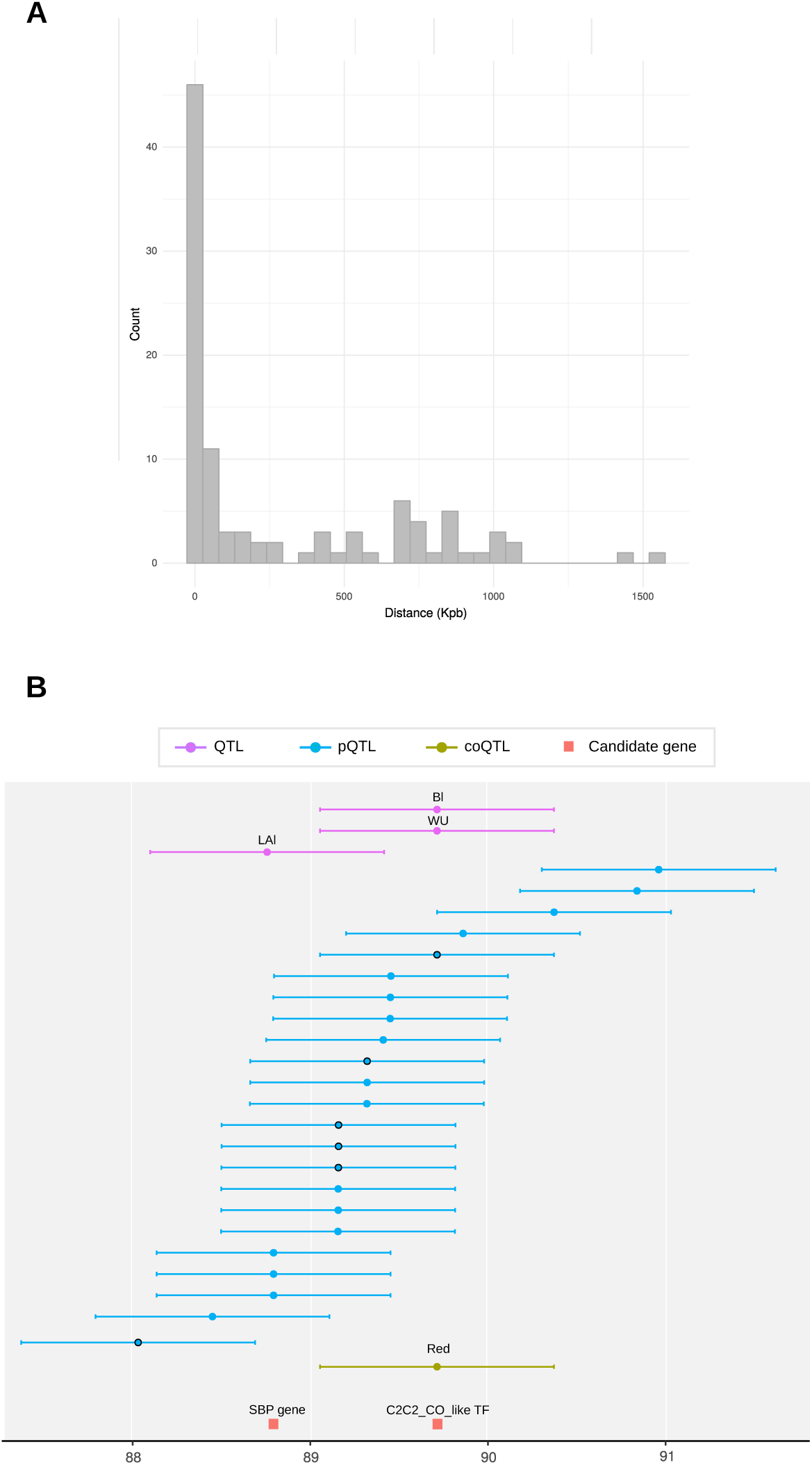
Identification of genomic regions involved in multi-scale genetic control. (A) Distribution of the distances between co-localizing QTLs and pQTLs. (B) Detailed view of the QTL, pQTL, coQTL detected in the region covered by the Hs52d hotspot on chromosome 5. Dots represent the SNPs determining the position of the QTLs and horizontal bars represent the linkage disequilibrium-based window around each SNP. Black circles indicate the pQTLs that co-localize with QTLs or coQTLs with a high correlations between the protein abundance and the phenotypic trait value or the module eigengene. The position of two transcription factors (an SBP gene, GRMZM2G111136, and a C2C2-CO-like transcription factor, GRMZM2G148772) representing promising candidate genes are indicated.

We further identified 11 pairs of SNPs corresponding to QTL/coQTL co-localizations, all in the WD condition (Supplemental Table S10). They involved three phenotypic traits (WU, Bl, LAl) and two co-expression modules including the WD-specific module (Fig. 5). These two modules were significantly enriched in stress-response proteins and in proteins involved in hormone metabolim and in reactive oxygen species detoxification (Supplemental Table S11). Ten of the 11 QTLs co-localizing with coQTLs also co-localized with pQTLs. The remaining QTL actually also co-localized with pQTLs, but with a low correlation between the phenotypic trait values and the protein abundance levels (|*r*_*corrected*_| < 0.23; Supplemental Table S10). By contrast, the correlation between trait values and eigengene was much higher (|*r*_*corrected*_| = 0.51), which indicates that proteins were more strongly related to ecophysiological traits when taken collectively through a co-expression module rather than taken individually.

Taken together, these results highlight the presence of loci associated with traits at different biological scales. In the WD condition, several of these loci showed multiple associations both at the proteome and the phenotype level (Fig. 5). On Chromosome 1, a locus spanning 33 kbp contained a QTL for LAl determined by SNP S1_5382845 as well as a coQTL for the green module and seven pQTLs, all determined by SNP AX-91427638. On Chromosome 5, a locus spanning 1.8 Mb between SNPs AX-91657926 and AX-91658235, contained three QTLs for LAl, Bl and WU, one coQTLs for the WD-specific module and six pQTLs. This region also contained hotspot Hs52d. On Chromosome 7, a single SNP (S7_162671160) determined the positions of two QTLs for LAl and WU, two coQTLs and seven pQTLs. On Chromosome 10, a locus spanning 1.3 Mb between SNPs S10_122802154 and S10_124095144, contained one QTL for LAl, one coQTL and eight pQTLs. This region also contained hotspot Hs103d. Note that in the WD condition, leaf area (LAl) was repeatedly associated with the green module (on Chromosomes 1, 7, 9, 10) and to proteins belonging to this module. Several of them were detoxification enzymes (*i.e.*, a putative polyphenol oxydase, GRMZM5G851266; two peroxydases, GRMZM2G085967 and GRMZM2G108153; a superoxide dismutase GRMZM2G025992; a glyoxalase GRMZM2G704005).

### Identification of candidate genes potentially involved in drought tolerance

Assuming that the genetic polymorphisms associated with protein abundance variations are within genes, we retrieved a list of one to 49 candidate genes for each of the 69 pairs of SNPs corresponding to a QTL/pQTL or QTL/coQTL co-localization (Supplemental Table S12). Based on gene annotation and the literature, we identified two particularly interesting cases.

First, on Chromosome 7, the SNP S7_162671160 was located in *ZmGH3.8* (GRMZM2G053338), which was the only candidate gene. *ZmGH3.8* is involved in indole-3-acetyl-amide conjugate biosynthesis. In agreement with the role of this gene in drought response (Feng et al. 2015), S7_162671160 was associated with the WD-specific module, WU and LAl and five stress-response proteins (endochitinase GRMZM2G051943, beta-D-glucanase GRMZM2G073079, peroxidase GRMZM2G085967, polyphenol oxydase GRMZM5G851266 and phospholipase D GRMZM2G061969).

Second, 14 candidate genes were identified in the region of Chromosome 5 covered by hotspot Hs52d, of which two could be associated with the expression variation of a high number of genes. One is a squamosa promoter-binding (SBP) gene (GRMZM2G111136) that is inducible by various abiotic stresses including drought (Mao et al. 2016). The other, a C2C2-CO-like transcription factor (GRMZM2G148772), was found to be significantly induced by drought and salinity stress in B73 leaves (Forestan et al. 2016). Hotspot Hs52d covered a region of *ca* 4 Mb in which we detected 26 pQTLs (many of which were located between the SBP gene and the C2C2-CO-like transcription factor), two coQTLs and four QTLs (Fig. 6B). A single SNP, AX-91658235 located only one kbp from the C2C2-CO-like transcription factor, determined the position of two QTLs, two pQTLs and one coQTL. Furthermore, SNP S5_88793314, located within the coding sequence of the SBP gene, determined the position of a QTL and a pQTL. Based on these results, we can hypothesize that hotspot Hs52d may correspond to two trans-acting factors for which the SBP gene and the C2C2-CO-like transcription factor represent good candidates.

## DISCUSSION

To better understand the molecular mechanisms associated with the genetic polymorphisms underlying the variations in ecophysiological traits related to drought tolerance, we used a proteomics-based systems genetics approach which allowed us to map 22,664 pQTLs at high-resolution. By relating pQTLs to protein functions and heritability, we showed that the level of genetic control over protein abundances depends on protein function. Notably, proteins involved in translation and energy metabolism exhibited few pQTLs, with a lack of local pQTLs and low heritability. As these two functional categories mainly contain ancient and evolutionarily conserved proteins (Goldman et al. 2010; Nelson and Junge 2015), our results suggest that evolutionarily ancient proteins have more constrained expressions and fewer associated pQTLs (Mähler et al. 2017; Popadin et al. 2014; Zhang and Yang 2015). They also support the recent hypothesis of Mähler et al. (2017) that, for genes experiencing reduced rates of molecular evolution, purifying selection on individual SNPs is associated with stabilizing selection on gene expression.

pQTLs were found throughout the genome but some of them clustered into hotspots, suggesting the presence of loci with pleiotropic effects on the proteome. The detection of QTL hotspots is highly dependent on the number of traits studied, the mapping resolution and the method used to cluster QTLs. This may explain why previous studies have reported hotspots ranging from hundreds of eQTLs (Munkvold et al. 2013; Christie et al. 2017; Orozco et al. 2012) to only a few tens of eQTL or pQTLs (Foss et al. 2011; Ghazalpour et al. 2011; Albert et al. 2014) or even no hotspot at all (Mähler et al. 2017). In our study, false hotspot detection was limited by having a high mapping resolution and by using a pQTL clustering method that takes into account LD variations across the genome (Negro et al. 2019). Based on co-localization with coQTLs, we ultimately cross-validated 15 condition-specific hotspots, suggesting that loci with pleiotropic effects on the proteome can interact with the environment.

By analyzing a diversity panel of 254 genotypes, we showed that many small changes in protein abundance, detected as significant because they occurred in a high number of genotypes, contributed to extensively remodel the proteome in water deficit conditions. In total, approximately 75% of quantified proteins responded significantly to environmental change. Up- and down regulated proteins were well differentiated in terms of function, and indicated that the photosynthetic, transcriptional and translational machineries were slowed down while stress responses and signalization mechanisms were activated. All these changes showed that plants clearly perceived a lack of water and presented a coordinated proteome response to water deficit.

Changes in abundance occurring in response to water deficit were associated with changes in the structure of the co-expression network. Indeed, we identified condition-specific modules, one of which, in the WD condition, was significantly enriched for stress-response proteins. Similarly, Munkvold et al. (2013) observed condition-specific modules related to biological processes in response to particular environmental conditions. Such modules suggest that, under environmental perturbation, sets of genes or proteins are collectively mobilized by condition-specific factors allowing plant cells to adapt. The WD-specific module was associated with several QTL/coQTL colocalizations and its eigengene was highly correlated with biomass, water use and leaf area. Although the approach used here is correlative, these results suggest that, under water deficit, stress-response proteins contribute to phenotypic responses, which is consistent with the fact that many QTL/pQTL co-localizations involved these types of proteins. One coQTL for the WD-specific module was located in a region of Chromosome 5 that also cumulated several QTLs, pQTLs and the hotspot Hs52d. This indicates that the co-expression observed for stress response proteins may be driven by condition-specific factors, the pleiotropic effects of which resonate across all layers of biological complexity up to phenotype. Altogether, these results suggest that an eigengene may be considered a more integrated molecular trait than protein abundance, and can help decipher the genotype-phenotype relationship by bridging the gap between the proteomic and phenotypic level.

Linking phenotypic variation to proteome variation revealed many QTL/pQTL co-localizations for which, using high mapping resolution, we identified a limited number of candidate genes.

Only two of the 69 QTLs detected in the WW condition, *vs* 39 of the 91 in the WD condition, co-localized with pQTLs. This difference could be explained by the hypothesis that under non-stress conditions, phenotypic variations are driven by many low contribution proteins, whose abundance is probably controlled by low effect genetic polymorphisms, whereas under water stress, phenotypic variations are mainly driven by stress response proteins under the genetic control of condition-specific factors. In agreement with this hypothesis, we robustly identified two genomic regions that could correspond to such factors. The first is located on Chromosome 7, where we identified *ZmGH3.8* as the sole candidate gene underlying two QTLs (for leaf area and water use), seven pQTLs, of which five were associated with proteins involved in stress responses, and two coQTLs, one of which was associated with the WD-specific module. In maize shoots, Feng et al. (2015) showed that the expression of *ZmGH3.8* was induced by auxin and reduced under polyethylene glycol treatment. The second region is located on Chromosome 5, in the region of the Hs52d hotspot, where we identified an SBP gene (GRMZM2G111136) and a C2C2-CO-like gene (GRMZM2G148772) as candidate genes underlying four QTLs, six pQTLs and one coQTLs. These two transcription factors have been previously shown to be induced by drought in maize (Mao et al. 2016; Forestan et al. 2016). In addition, SBP genes constitute a functionally diverse family of transcription factors involved in plant growth and development (Preston and Hileman 2013). Due to their potential implication in GxE interactions and because of their roles both in plant growth and development and in drought response, Z*mWRKY48, ZmGH3.8*, SBP and C2C2-CO-like genes represent promising candidates for drought tolerance breeding.

To conclude, our systems genetics approach which incorporates MS-based proteomics data has yielded several new results regarding the drought response in maize. First, we point out that the strength of the genetic control over protein abundance is related to protein function and also probably to the evolutionary constraints on protein expression. Then, we show that even mild water deficit strongly remodels the proteome and induces a reprogramming of the genetic control of the abundance of many proteins and notably those involved in stress responses. QTL/pQTL co-localizations are mostly found in the WD condition indicating that this reprogramming also affects the phenotypic level. Finally, we identify candidate genes that are potentially responsible for both the co-expression of stress-response proteins and the variation of ecophysiological traits under water deficit. Taken together, our findings provide novel insights into the molecular mechanisms of drought tolerance and suggest some pathways for further research and breeding. Our study also demonstrates that proteomics has now reached enough maturity to be fully exploited in systems studies necessitating large-scale experiments.

## METHODS

### Plant material and experiment

Plant material and growth conditions are described in full details in Prado et al. (2018) and in the Supplemental Methods. In brief, a diversity panel of maize hybrids was obtained by crossing a common flint parent (UH007, paternal parent) with 254 dent lines. Two levels of soil water content were applied: well-watered (soil water potential of −0.05 MPa) and water deficit (soil water potential of -0.45 MPa). Hybrids were replicated three times in each watering condition. Leaf sampling was performed at the pre-flowering stage in two replicates per hybrid and water condition.

### Protein extraction and digestion

Protein extraction and digestion procedures are described in full detail in the Supplemental Methods. In brief, proteins were extracted from frozen ground leaf samples using a standard protocol for protein precipitation with trichloroacetic acid and acetone solution. Tryptic digestion was performed after solubilization, reduction and alkylation of the proteins. The resulting peptides were desalted by solid phase extraction using polymeric C18 columns.

### LC-MS/MS analyses

Samples were analyzed by LC-MS/MS in batches of 96. Analyses were performed using a NanoLC-Ultra System (nano2DUltra, Eksigent, Les Ulis, France) connected to a Q-Exactive mass spectrometer (Thermo Electron, Waltham, MA, USA). A 400 ng protein digest was loaded at 7.5 μl.min^−1^ on a Biosphere C18 pre-column (0.3 × 5 mm, 100 Å, 5 μm; Nanoseparation, Nieuwkoop, Netherlands) and desalted with 0.1% formic acid and 2% ACN. After 3 min, the pre-column was connected to a Biosphere C18 nanocolumn (0.075 × 150 mm, 100 Å, 3 μm, Nanoseparation). Buffers were 0.1% formic acid in water (A) and 0.1% formic acid and 100% ACN (B). Peptides were separated using a linear gradient from 5 to 35% buffer B for 40 min at 300 nl.min^−1^. One run took 60 min, including the regeneration step at 95% buffer B and the equilibration step at 95% buffer A. Ionization was performed with a 1.4-kV spray voltage applied to an uncoated capillary probe (10 μm tip inner diameter; New Objective, Woburn, MA, USA). Peptide ions were analyzed using Xcalibur 2.2 (Thermo Electron) in a data-dependent acquisition mode as described in the Supplemental Methods.

### Peptide and protein identification

Peptide identification was performed using the MaizeSequence genome database (Release 5a, 136,770 entries, https://ftp.maizegdb.org/MaizeGDB/FTP/) supplemented with 1,821 French maize inbred line F2 sequences with present/absent variants (PAVs) (Darracq et al. 2018) and a custom database containing standard contaminants. Database searches were performed using X!Tandem (Craig and Beavis 2004) (version 2015.04.01.1) and protein inference was performed using a homemade C++ version of X!TandemPipeline (Langella et al. 2017) specifically designed to handle hundreds of MS run files. Parameters for peptide identification and protein inference are described in the Supplemental Methods. The false discovery rate (FDR) was estimated at 0.06% for peptides and 0.04% for proteins.

Functional annotation of proteins was based on MapMan mapping (Thimm et al. 2004; Usadel et al. 2009) (Zm_B73_5b_FGS_cds_2012 available at https://mapman.gabipd.org/) and on a custom KEGG classification built by manually attributing the MapMan bins to KEGG pathways (Dillmann, pers. com.).

### Peptide and protein quantification

Peptide quantification was performed using MassChroQ version 2.1.0 (Valot et al. 2011) based on extracted ion chromatograms (XIC) with the parameters described in the Supplemental Methods. Peptide quantification data were filtered to remove genotypes represented by only one or two samples instead of the expected four, as well as outlier samples for which we suspected technical problems during sample preparation or MS analysis. In the end, the MS dataset included 977 samples.

Proteins were quantified from peptides using two complementary methods. *i. XIC-based quantification:* Proteins were quantified based on peptide intensity data filtered and normalized as described in the Supplemental Methods. We excluded proteins that were quantified by only one peptide. As samples were analyzed by LC-MS/MS in batches over a period of several months, we observed a strong batch effect on normalized peptide intensities. To correct this batch effect, we fitted a linear model to log-transformed intensity data and subtracted the component due to batch effects. Then, for each protein, we modeled the peptide data using a mixed-effects model derived from Blein-Nicolas et al. (2012) and described in the Supplemental Methods. Protein abundance was subsequently computed as adjusted means from the model’s estimates. *ii. Spectral counting (SC)-based quantification:* Proteins that could not be quantified with XIC because their peptides had too many missing intensity values were quantified based on their number of assigned spectra. Proteins with a spectral count < 2 in any of the samples were discarded. Normalization was then performed as described in the Supplemental Methods. As in XIC-based quantification, we corrected the batch effect by fitting a linear model to square-root transformed and normalized protein abundances. Analysis of variance (ANOVA) was subsequently performed using the mixed-effects model described in the Supplemental Methods.

### Genome wide association study

GWAS was performed on protein abundances estimated in each watering condition using the single locus mixed model described in Yu et al. (2006). The variance-covariance matrix was determined as described in Rincent et al. (2014) by a kinship matrix derived from all SNPs except those on the Chromosome containing the SNP being tested. SNP effects were estimated by generalized least squares and their significance was tested with an F-statistic. An SNP was considered significantly associated when -log_10_(*P-value*) > 5. A set of 961,971 SNPs obtained from line genotyping using a 50 K Infinium HD Illumina array (Ganal et al. 2011), a 600 K Axiom Affymetrix array (Unterseer et al. 2014) and a set of 500 K SNPs obtained by genotyping-by-sequencing (Negro et al. 2019) were tested. Analyses were performed with FaST-LMM (Lippert et al. 2011) v2.07. Only SNPs with minor allele frequencies > 5% were considered.

Inflation factors were computed as the slopes of the linear regressions on the QQplots between observed -log_10_(*P-value*) and expected -log_10_(*P-value*). Inflation factors were close to 1 (median of 1.08 and 1.06 in the XIC-based and SC-based sets, respectively), indicating low inflation of *P-values*.

### Detection of QTLs from significantly associated SNPs

Three different methods implemented in R (R core team 2013) version 3.3.3 were used to summarize significantly associated SNPs into pQTLs. *i. The genetic method:* two contiguous SNPs were considered to belong to a same QTL when the genetic distance separating them was less than 0.1 cM. *ii. The LD-based method:* two contiguous SNPs were considered to belong to a same QTL when their LD-based windows (Negro et al. 2019) overlapped. *iii. The geometric method:* for each chromosome, we ordered the SNPs according to their physical position. Then, we smoothed the -log_10_(*P-value*) signal by computing the maximum of the -log_10_(*P-values*) in a sliding window containing *N* consecutive SNPs. An association peak was detected when the smoothed -log_10_(*P-value*) signal exceeded a max threshold *M*. Two consecutive peaks were considered to be two different QTLs when the -log_10_(*P-value*) signal separating them dropped below a min threshold *m*. The parameters for QTL detection were fixed empirically at *N=*500, *M*=5 and *m=*4. For the three methods described above, the position of a QTL was determined by the SNP exhibiting the highest -log_10_(*P-value*). A pQTL was considered local if it was within 1 Mb upstream or downstream of the coding sequence of the gene encoding the corresponding protein.

### Complementary data analyses

The following complementary data analyses were performed with R (R core team 2013) version 3.3.3. *i. Broad sense heritability of protein abundance:* For each protein, the broad sense heritability of abundance was computed for each of the two watering conditions from a mixed-effects model as described in the Supplemental Methods. *ii. detection of pQTL hotspots:* for each SNP position, we counted the number of pQTLs (*N*) located within its LD-based window (Negro et al. 2019). The threshold used to detect a hotspot was set at the 97% quantile of the distribution of *N. iii. Protein co-expression analysis:* Protein co-expression analysis was performed using the WGCNA R package (Langfelder and Horvath 2008) with the parameters described in the Supplemental Methods. Using a procedure developed to correct the bias due to population structure and/or relatedness in the LD measure and implemented in the LDcorSV R package (Mangin et al. 2012), we computed pair-wise Pearson’s correlations corrected by structure and kinship (|*r*_*corrected*_|) and used them as the input similarity matrix. Graphical representations of the resulting networks were performed with Cytoscape (Shannon et al. 2003) v3.5.1 using an unweighted spring embedded layout. *iv. QTL co-localization:* We considered QTLs to co-localize when they meet the following two criteria. First, the LD-based windows around the QTLs (Negro et al. 2019) should overlap. Second, the absolute value of the Pearson’s correlation of coefficient corrected by structure and kinship (the |*r*_*corrected*_| mentioned above) between the values of the ecophysiological traits associated with the QTLs should be greater than 0.3. We determined this value empirically, in the absence of a statistical test to test the significance of the corrected correlation. *v. Candidate gene identification:* For each QTL/pQTL co-localization, gene accessions found within the interval defined by the intersection between the LD-based windows around the QTL and the pQTL were retrieved from the MaizeSequence genome database (Release 5a). Low confidence gene models and transposable elements were not considered.

## DATA ACCESS

The raw MS output files were deposited online using PROTICdb (Langella et al. 2007; Ferry-Dumazet et al. 2005; Langella et al. 2013) at the following URL: http://moulon.inra.fr/protic/amaizing (DOI 10.15454/1.5736519296148652E12, currently available with the following username: reviewer and password: reviewer) and at MassIVE at the following URL: ftp://MSV000084619@massive.ucsd.edu (<u>doi:10.25345/C5Q684</u>). They will be made freely available after publication. Detailed information on all peptides and proteins identified in the LC-MS/MS runs as well as peptide intensities and protein abundances obtained for each sample are also freely available on PROTICdb at the same URL.

Phenotypic data are available online using the PHIS information system (Neveu et al. 2019) at the following URL: http://www.phis.inra.fr/openphis/web/index.php?r=project%2Fview&id=Systems+genetics+for+maize+drought+tolerance+%28Amaizing+project%29. Earlyeaf area (LAe) was defined at the seven leaves stage, representing 24 d_20°C_ (thermal time in equivalent days at 20°C). Late leaf area (LAl) was defined at the 12 leaves stage, representing 45 d_20°C_.

Genotyping data are available at the following URL: https://doi.org/10.15454/GAHEU0

## ACKNOWLEDGMENTS

This work was supported by the Agence Nationale de la Recherche project ANT-10-BTBR-01 (Amaizing). Proteomics analyses were performed on the PAPPSO platform (http://pappso.inra.fr) which is supported by INRA (http://www.inra.fr), the Ile-de-France regional council (https://www.iledefrance.fr/education-recherche), IBiSA (https://www.ibisa.net) and Saclay Plant Sciences-SPS (ANR-17-EUR-0007). The authors want to thank Sylvie Coursol for her critical review of the manuscript, Hélène Corti for her help in sample preparation and Olivier Langella for having specially developed a pipeline to upload the proteomics data on ProticDB. They are also grateful to people from INRA LEPSE: François Tardieu for his contribution to the coordination of the plant experiment; Benoît Suard, Pauline Sidawi and Olivier Martin for their technical assistance during the experiment; Santiago Alvarez Prado for his contribution to plant traits and QTL analysis.

## AUTHORS CONTRIBUTION

AC and MZ designed the research; CW designed and coordinated the plant experiment in PhenoArch and the genetic analysis of plant traits; LBC performed the plant experiment and analyzed the image-based phenotypic data; MBN and TB performed the proteomics experiments; SSN developed the GWAS pipeline and performed the genotyping quality control; SDN performed the genotyping and estimated local LD; MBN and MZ analyzed the proteomics data, MBN performed the systems genetics study and wrote the manuscript. All authors discussed the results and read and approved the final manuscript.

## DISCLOSURE DECLARATION

The authors declare no competing interest.

## Notes

### Competing Interest Statement

The authors have declared no competing interest.

### Summary of Updates

The title has been changed and peak counting data have been replaced by spectral counting data

