## supplemental figures for "A systems genetics approach reveals environment-dependent associations between SNPs, protein co-expression and drought-related traits in maize"

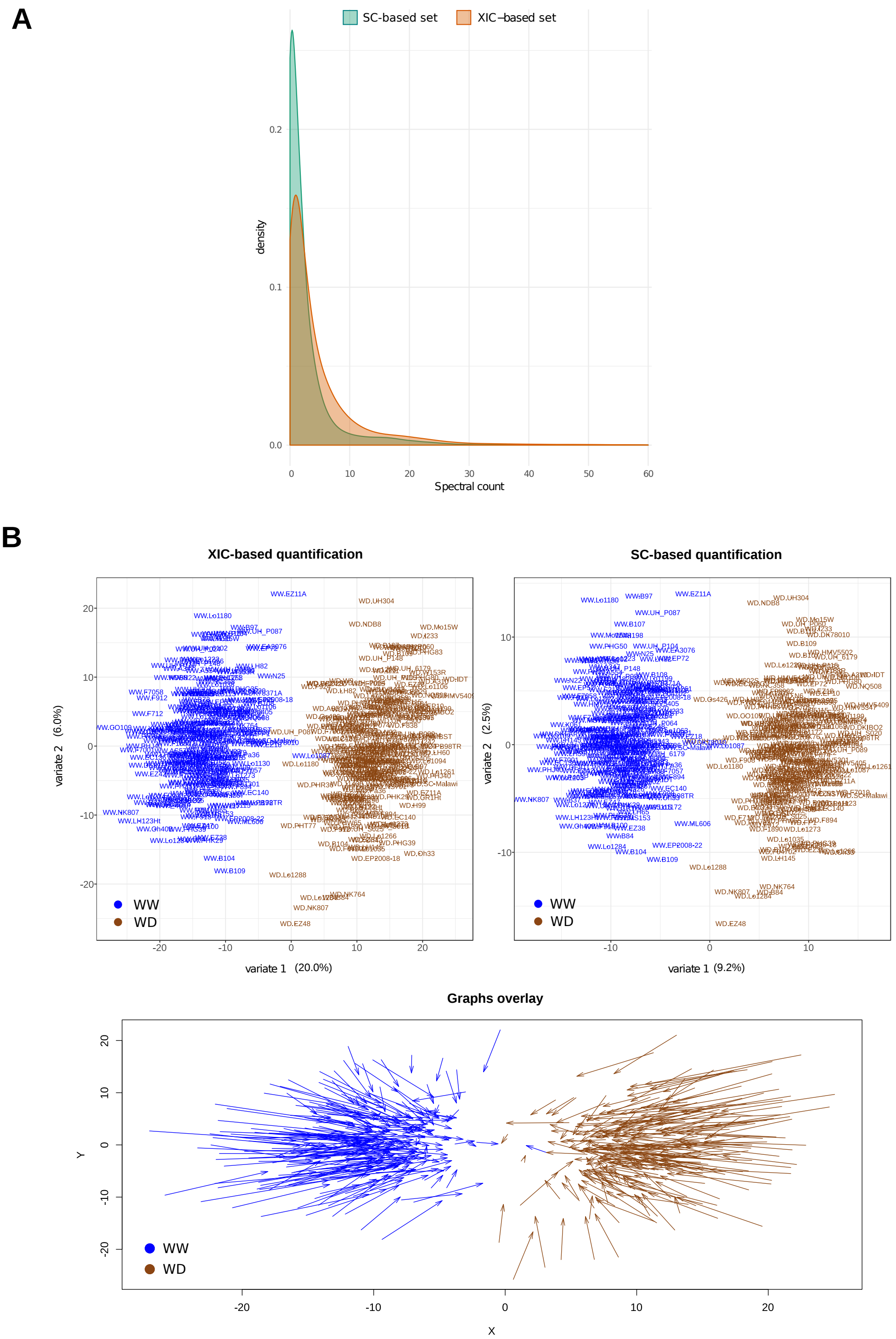


**Supplemental Figure S1. The XIC-based and SC-based protein sets exhibit different characteristics.** (A) Distribution of the spectral counts in the two sets. (B) Graphical representations of a partial least squares regression relating abundances estimated from spectral counts to abundances estimated from peptide intensities. Here, only proteins from the XIC-based set were used. They were quantified by using the two methods (XIC-based and SC-based) in order to compare the discrimination power provided by each method. The analysis was performed using the Mixomics R package. Top: samples representation in the space spanned by the first two components for each quantification method. Bottom: superposition of the samples representation obtained for the two quantification methods. Each sample is indicated by an arrow whose start indicates the location of the sample when quantification is based on XIC and tip indicates the location of the sample when quantification is based on SC. Orientation of the arrows indicates that the two watering conditions are better separated when proteins are quantified based on XIC.

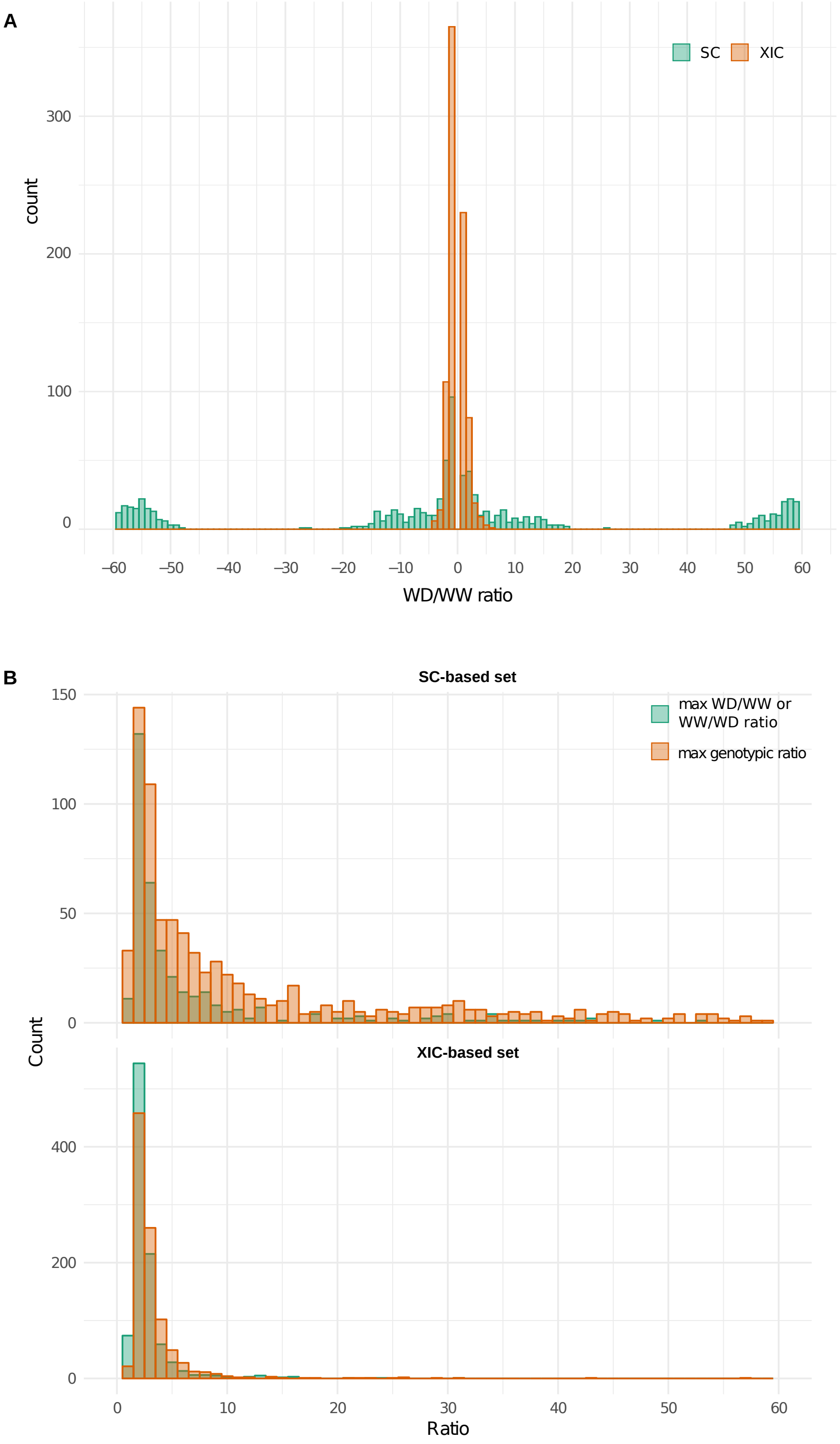


**Supplemental Figure S2. Distribution of maximum amplitude of abundance variation in response to water deficit or to a genotypic change.** (A) Distribution of the maximum WD/WW ratio for proteins showing significant abundance change in response to water deficit. (B) Comparison of the distributions of maximum amplitude of abundance variation in response to water deficit and genotypic change. In this case, the maximum amplitude of abundance variation in response to water deficit was the highest WD/WW or WW/WD ratio found across all genotypes. To compute the maximum amplitude of abundance variation in response to a genotypic change over the two conditions, we first computed the average protein abundance in each genotype and then the ratio between the highest and the lowest average abundances.

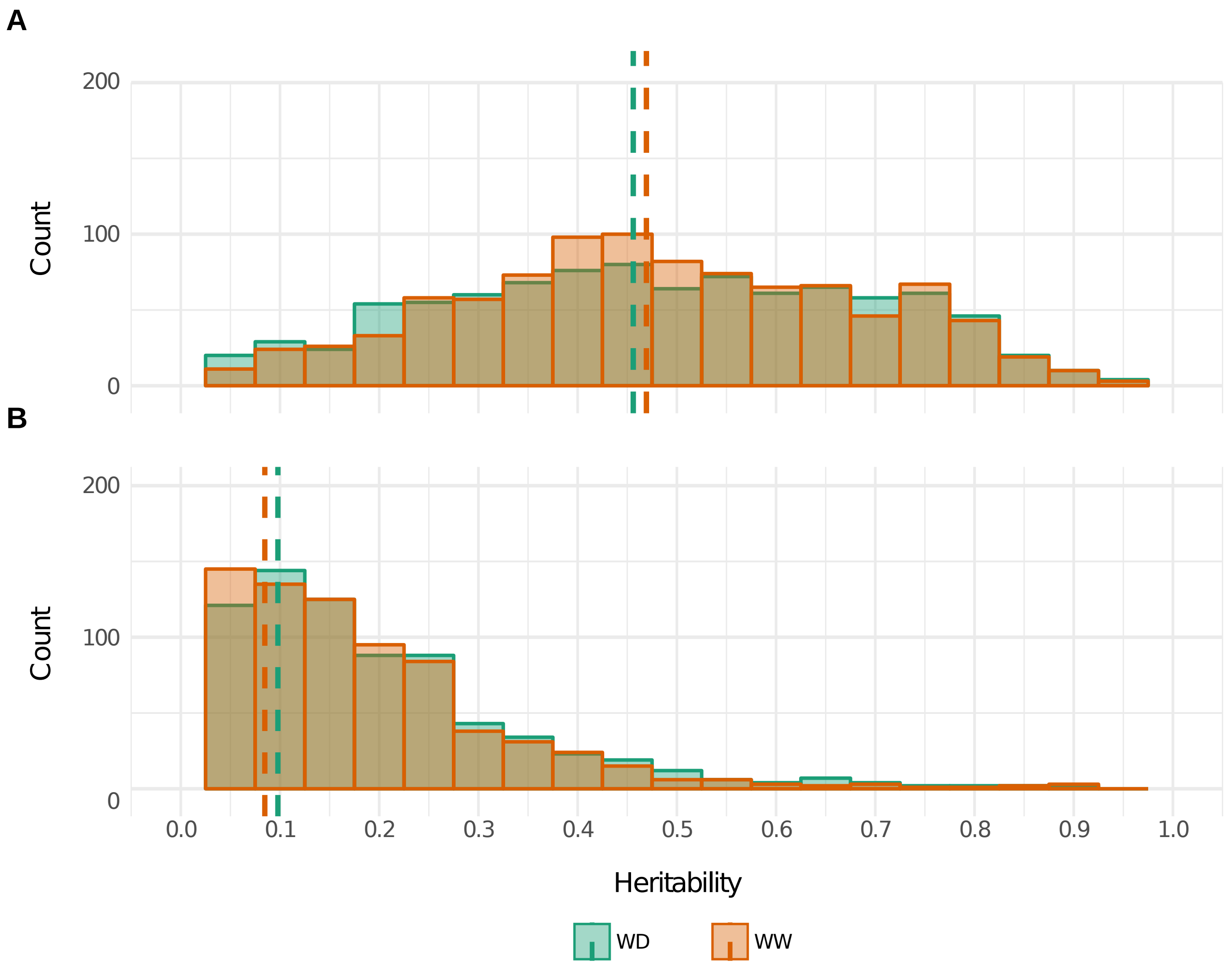


**Supplemental Figure S3. Distributions of broad sense heritability for protein abundances in the WW and WD conditions.** (A) in the XIC-based set. (B) in the SC-based set. Dashed lines indicate the median values.

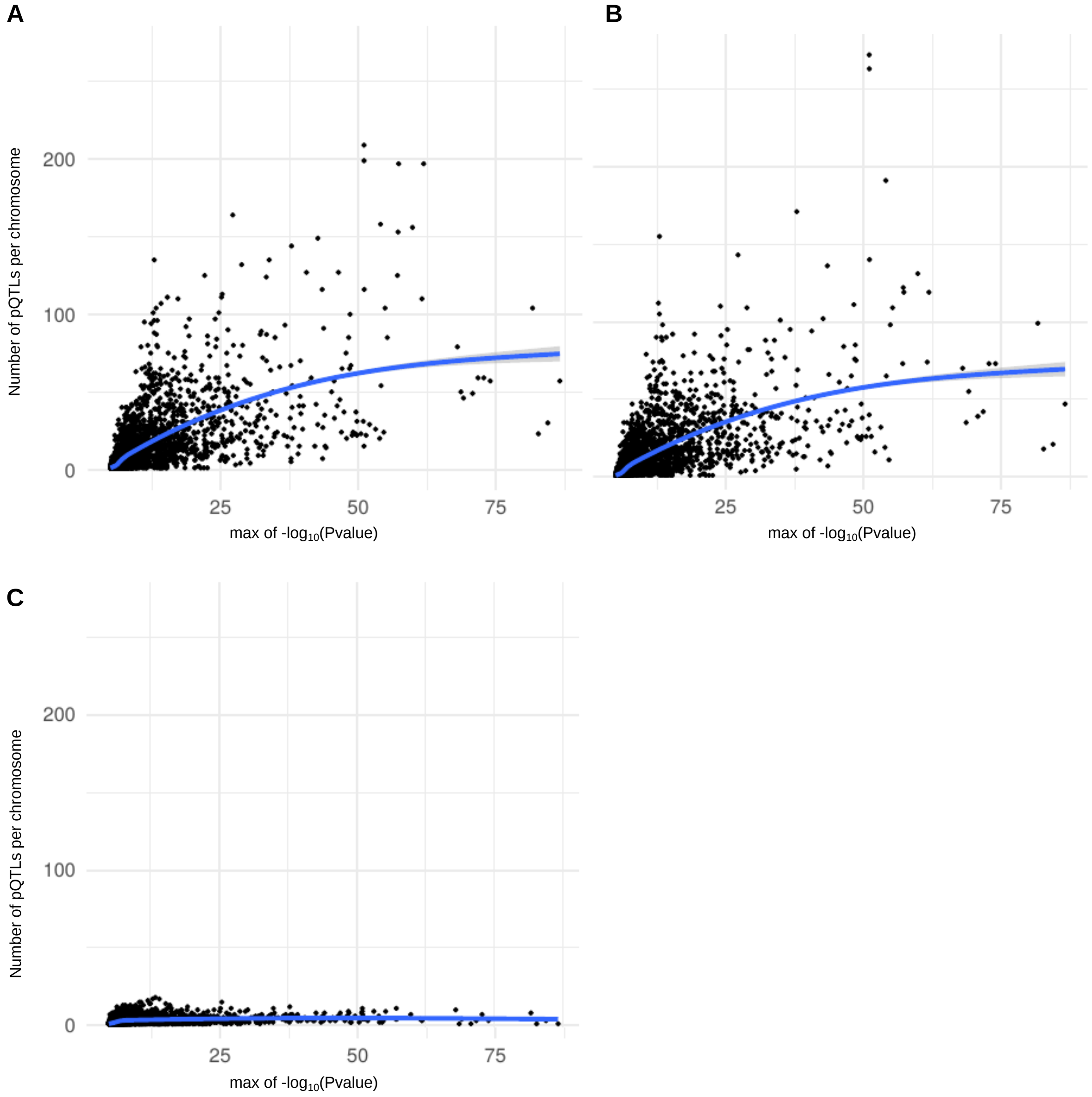


**Supplemental Figure S4. Effect of the methods of detection of association peaks on the number of pQTLs detected.** Each graph represents the relationships between the number of pQTLs detected per chromosome and the *P-value* of the most strongly associated pQTL of the corresponding chromosome. (A) LD-based method, (B) Genetic method with a 0.1 cM window, (C) Geometric method (see Materials & Methods for details).

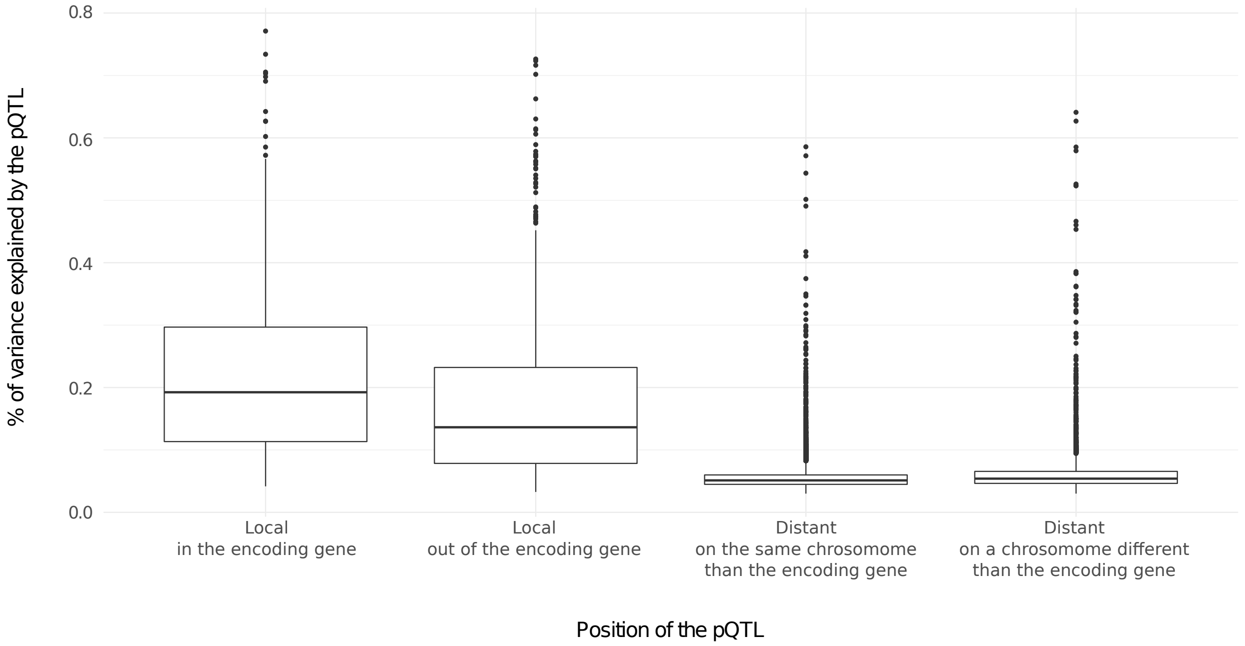


**Supplemental Figure S5. Boxplot of the percentage of variance explained by a pQTL depending on its position relatively to the gene encoding the protein to which it is associated.**

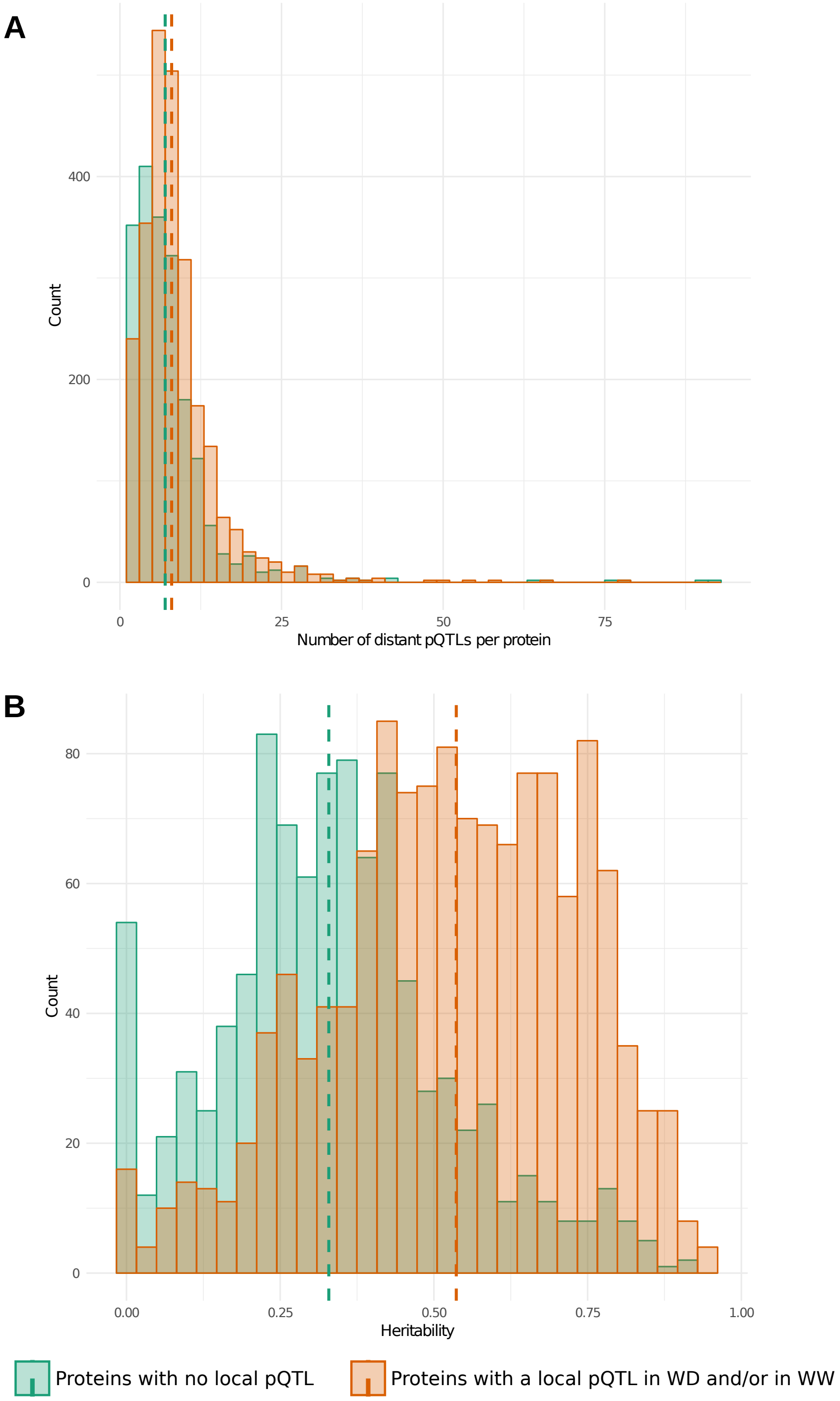


**Supplemental Figure S6. Caracterization of the proteins for which no local pQTL was detected.** (A) Distributions of the number of distant pQTLs per protein in the set of proteins for which no local pQTL was detected *versus* the set of proteins for which a local pQTL was detected in at least one condition. (B) Distributions of heritability in the set of proteins for which no local pQTL was detected *versus* the set of proteins for which a local pQTL was detected in at least one condition. Dashed lines represent median values.

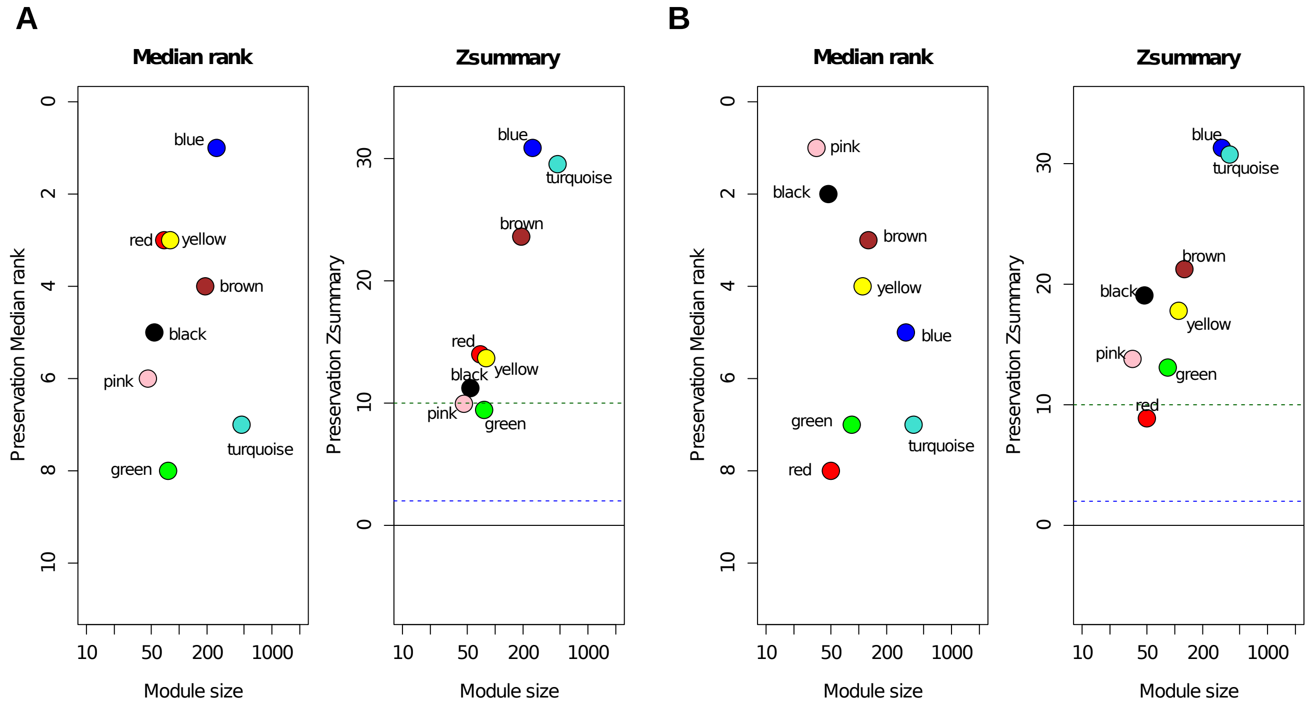


C

| Consensus modules (728 proteins) | WW modules (1,240 proteins) | | | | | | | | WD modules (1,165 proteins) | | | | | | | |
|---|---|---|---|---|---|---|---|---|---|---|---|---|---|---|---|---|
| | 46 | 80 | 471 | 54 | 69 | 253 | 76 | 191 | 35 | 84 | 390 | 47 | 322 | 110 | 127 | 50 |
| 209 | 1 | 1 | 5 | 2 | 2 | 185 | 6 | 2 | 0 | 1 | 190 | 1 | 3 | 0 | 5 | 4 |
| 26 | 0 | 0 | 1 | 0 | 25 | 0 | 0 | 0 | 0 | 0 | 4 | 21 | 1 | 0 | 0 | 0 |
| 18 | 0 | 0 | 4 | 0 | 0 | 1 | 0 | 4 | 0 | 18 | 0 | 0 | 0 | 0 | 0 | 0 |
| 78 | 0 | 2 | 1 | 0 | 1 | 1 | 1 | 71 | 0 | 1 | 12 | 0 | 1 | 3 | 58 | 3 |
| 40 | 0 | 1 | 8 | 2 | 0 | 2 | 0 | 26 | 1 | 6 | 5 | 0 | 3 | 0 | 24 | 0 |
| 15 | 0 | 0 | 7 | 3 | 0 | 0 | 0 | 5 | 0 | 0 | 2 | 0 | 3 | 10 | 0 | 0 |
| 52 | 0 | 42 | 2 | 0 | 0 | 0 | 0 | 8 | 0 | 0 | 3 | 0 | 2 | 46 | 0 | 0 |
| 18 | 0 | 4 | 11 | 0 | 0 | 0 | 0 | 3 | 0 | 14 | 0 | 0 | 0 | 2 | 0 | 2 |
| 20 | 0 | 0 | 0 | 20 | 0 | 0 | 0 | 0 | 0 | 2 | 2 | 0 | 2 | 1 | 1 | 0 |
| 12 | 0 | 0 | 0 | 0 | 0 | 0 | 0 | 0 | 0 | 0 | 0 | 0 | 2 | 0 | 0 | 0 |
| 211 | 0 | 2 | 183 | 1 | 12 | 3 | 5 | 4 | 2 | 2 | 4 | 8 | 186 | 5 | 0 | 0 |
| 29 | 1 | 0 | 26 | 0 | 0 | 0 | 1 | 0 | 0 | 2 | 17 | 3 | 3 | 0 | 0 | 0 |
| | 44 | 28 | 223 | 26 | 29 | 61 | 63 | 68 | 32 | 38 | 151 | 14 | 116 | 43 | 39 | 41 |

**Supplemental Figure S7. Similarities among the WW and WD networks.** (A) Preservation of WW modules in WD data. Zsummary is a composite measure summaring the Z scores computed on several preservation statistics. Zsummary < 2 (blue dashed line) indicates no preservation; 2 < Zsummary < 10 indicates weak to moderate preservation; Zsummary > 10 indicates good preservation. Median rank is based on the ranks of the observed preservation statistics. It does not show dependence on the module size. (B) Preservation of WD modules in WW data. (C) Correspondence of the WW and WD modules (in column) with consensus modules (in row) that contain proteins co-expressed in the two conditions. Numbers above and on the left of the tables indicate the total number of proteins in each module. Numbers in the tables indicate the number of proteins in the intersection of a WW or WD module with a consensus module. The numbers of proteins of the WW and WD modules that were not mapped to a consensus module are shaded in grey at the bottom of the tables. Signifiance of the Fisher's exact test for the overlap of two modules is color-coded with the scale of $\log_{10}(1/P\text{-}value)$ shown on the right. No correspondence with a consensus module was found for the pink and green modules in the WW condition nor for the pink and red modules in the WD condition, indicating that these two modules are condition specific.

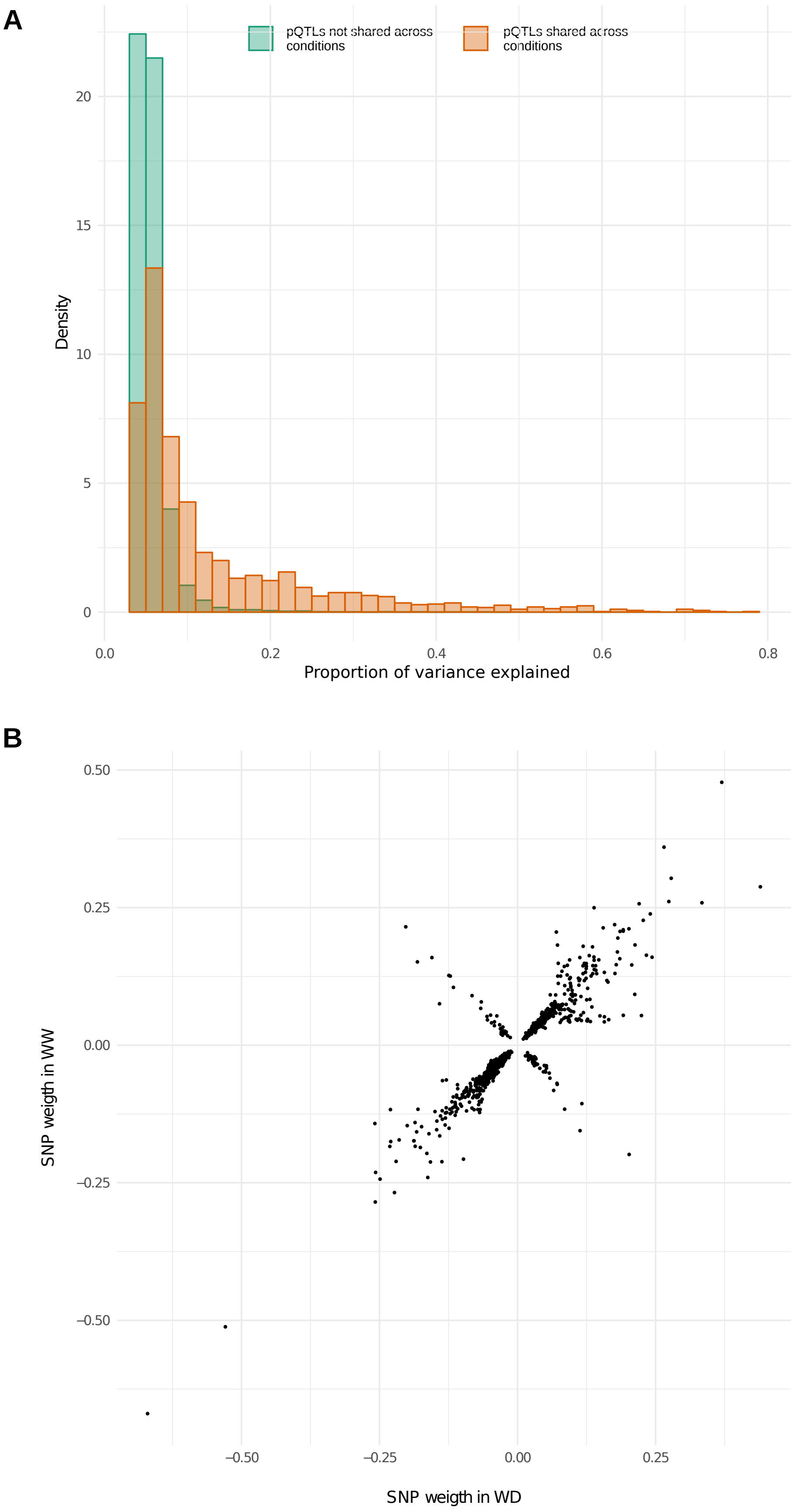


**Supplemental Figure S8. pQTLs shared across conditions.** (A) Distributions of the proportion of variance explained by the pQTLs shared and not shared across the two watering conditions. (B) Effects of the pQTLs shared across conditions on the abundances of the proteins they were associated to**.**
