## supplemental material for "A systems genetics approach reveals environment-dependent associations between SNPs, protein co-expression and drought-related traits in maize"

######################################################
# This is the R script used to filter and normalize peptide intensity data
######################################################


############# load libraries
	library(made4)


############# Prepare data

#!!!!!!!!!!! Note that Masschroq output file containing peptide intensities was too big to be loaded in R. It was split into 3 subfiles using a Python script: 
		# one containing the peptide intensities "XIC_result_130416_pep_quanti.tsv"
		# one containing the peptide retention times "XIC_result_130416_pep_rt.tsv"
		# one containing the peptide sequences "XIC_result_130416_pep_sequence.tsv"

	### load metadata
	setwd("my_repository")
	metadata=read.table("metadata.tsv", header=TRUE, sep="\t", stringsAsFactors=FALSE) #1,103 obs.

	### load peptide intensities
	tab.quanti=read.table("XIC_result_130416_pep_quanti.tsv", header=TRUE, sep="\t", dec=".", stringsAsFactors=FALSE)  #22,474,469 obs
	colnames(tab.quanti)=c("msrun_ID", "peptiz", "logQ")
	indice=match(tab.quanti$msrun_ID, metadata$msrun_ID)
	tab.quanti$hybrid_ID=metadata$hybrid_ID[indice]
	tab.quanti$watering_condition=metadata$watering_condition[indice]
	tab.quanti$replicate=metadata$replicate[indice]
	tab.quanti$order_of_sample_analysis=metadata$order_of_sample_analysis[indice]
	tab.quanti$plate_number=metadata$plate_number[indice]
	tab.quanti$sample_ID=metadata$sample_ID[indice]
	tab.quanti$sample_ID=as.character(tab.quanti$sample_ID)

	### load peptide retention times
	tab.rt=read.table("XIC_result_130416_pep_rt.tsv", header=TRUE, sep="\t", dec=".", stringsAsFactors=FALSE) #22,474,469 obs
	colnames(tab.rt)=c("msrun_ID", "peptiz", "rt", "rtdiff")

	### load peptide sequences
	tab.seq=read.table("XIC_result_130416_pep_sequence.tsv", header=TRUE, sep="\t", dec=".", stringsAsFactors=FALSE) #22,474,469 obs
	tab.seq$peptiz=paste(tab.seq$peptide, tab.seq$z, sep="-")
	tab.seq$msrun_ID=tab.rt$msrun_ID

	### load peptide mass modifications due to single amino acid polymorphism (SAP)
	tab.mut=read.table("list_total_peptide_pour_isotopes140416.tsv", header=TRUE, sep="\t", dec=".", stringsAsFactors=FALSE) #34,336 obs

	### load identified proteins and peptides
	tab.prot=read.table("proteins130416_corrected.tsv", header=TRUE, sep="\t", dec=".",stringsAsFactors=FALSE)  #39,813 obs
	colnames(tab.prot)[2]="sousousgroupe"
	spli=strsplit(tab.prot$sousousgroupe, ".", fixed=TRUE)
	protein=NULL
	for (i in (1:nrow(tab.prot))){
		protein=c(protein, paste(spli[[i]][1], spli[[i]][2], sep="."))
	}
	tab.prot$protein=protein
	tab.prot=tab.prot[,-2]
	tab.prot=unique(tab.prot[,c("peptide", "protein")])


############# Filters on samples

	### remove 12 plants sampled by error (genotypes that were grown in the greenhouse at the same time but that were not part of the experiment)
	count=as.data.frame(table(metadata$hybrid_ID))
	to_remove=count[count$Freq<3,1] 
	tab.quanti=tab.quanti[!tab.quanti$hybrid_ID %in% to_remove,] 

	### remove technical replicates analyzed on plate 12 (we decided to keep those analyzed on plate 11 because they had globally more quantified peptides)
	count=table(metadata$sample_ID)
	replicates=count[count>1]
	tab.quanti=tab.quanti[!(tab.quanti$sample_ID %in% names(replicates) & tab.quanti$plate_number=="plaque12"),] 

	### remove dubious LC-MS/MS runs showing low numbers of quantified peptides
	tab.quanti=tab.quanti[!tab.quanti$sample_ID %in% c(232, 240, 870,619, 843),] 


############# Filters peptides showing highly variable RT

	### filter on peptides' retention times
	tab.rt=tab.rt[tab.rt$msrun_ID %in% unique(tab.quanti$msrun_ID),] 
	total_rt <-na.omit(aggregate(tab.rt$rt,list(peptiz = tab.rt$peptiz),FUN=sd))
	hist(total_rt$x,nclass=500,freq=T,xlim=c(0,40),xlab="Standard deviation (s)") # mode around 5 sec
	peptiz_with_good_rt = total_rt$peptiz[total_rt$x<15] 
	tab.quanti=tab.quanti[tab.quanti$peptiz %in% peptiz_with_good_rt,]


############# Filter outliers samples based on PCA representation
	tab.acp=as.data.frame(tapply (tab.quanti$logQ, list(tab.quanti$peptiz,tab.quanti$sample_ID), FUN=sum)) 
	tab.acp=na.omit(tab.acp) 

	pca=ord(tab.acp, type="pca", trans=TRUE)
	sm=sum(pca$ord$eig)
	pound = round((pca$ord$eig/sm*100),digits = 1)
	max(pca$ord$li[,1])->maxx
	min(pca$ord$li[,1])->minx
	max(pca$ord$li[,2])->maxy
	min(pca$ord$li[,2])->miny
	max(pca$ord$li[,3])->maxz
	min(pca$ord$li[,3])->minz

	pca$ord$li=merge(pca$ord$li, metadata, by.x="row.names", by.y="sample_ID")
	mycol=cbind.data.frame(col_plate=c("black", "red","green3","blue","cyan","magenta","yellow","gray","purple3", "pink", "darkorange", "lightsalmon2"), plate_number=unique(pca$ord$li$plate_number))
	pca$ord$li=merge(pca$ord$li,mycol, "plate_number")
	mycol=cbind.data.frame(col_condition=c("brown", "blue"), watering_condition=unique(pca$ord$li$watering_condition))
	pca$ord$li=merge(pca$ord$li,mycol, "watering_condition")

	plot(pca$ord$li[,c("Axis1","Axis2")],type="n",xlab=paste("Axe1(",pound[1],"%)",sep=" "),ylab=paste("Axe2(",pound[2],"%)",sep=" "), xlim=c(minx,maxx), ylim=c(miny,maxy), main="couleurs par numero de plaque")
	abline(h=0,v=0,col="grey",lty=2)
	text(pca$ord$li[,c("Axis1","Axis2")], as.character(pca$ord$li$Row.names), col=as.character(pca$ord$li$col_plate))
	# => samples 697,698,699,700,701,702,703,704,705,706,707,708 appear as outliers on the first plan of PCA. These 12 samples were located on a sample plate line. 
 
	plot(pca$ord$li[,c("Axis3","Axis4")],type="n",xlab=paste("Axe3(",pound[3],"%)",sep=" "),ylab=paste("Axe4(",pound[4],"%)",sep=" "), main="couleurs par traitement")
	abline(h=0,v=0,col="grey",lty=2)
	text(pca$ord$li[,c("Axis3","Axis4")], pca$ord$li$sample, col=as.character(pca$ord$li$col_condition)) # conditions are separated on axis 4	

	tab.quanti=tab.quanti[!tab.quanti$sample_ID %in% c(697,698,699,700,701,702,703,704,705,706,707,708),] #19946493 obs., 977 samples, 31283 peptidez


############# Intensity normalization 

	### computing the bias taking as reference the msrun that presents the highest number of quantified peptiz 
	msrun=unique(tab.quanti$msrun_ID)
	nbrun=length(msrun) #977

	count=table(tab.quanti$msrun) 
	ref.msrun=names(count[count==max(count)])

	mat.obj=tapply(tab.quanti$logQ,list(tab.quanti$peptiz,tab.quanti$msrun_ID),FUN=sum)
	ref=mat.obj[,ref.msrun]
	mat.ref=as.data.frame(matrix(rep(ref,nbrun),ncol=nbrun)) 
	tab.bias=mat.ref-mat.obj 
	colnames(tab.bias)=colnames(mat.obj)
	tab.bias=stack(tab.bias)
	bias.peptiz=rep(rownames(mat.obj),nbrun)
	tab.bias=cbind.data.frame(bias.peptiz, tab.bias)
	colnames(tab.bias)=c("peptiz","bias","msrun_ID")
	indice=match(paste(tab.bias$msrun_ID, tab.bias$peptiz), paste(tab.rt$msrun_ID, tab.rt$peptiz))
	tab.bias$rt=tab.rt$rt[indice]

	### normalisation by smoothing the bias
	tab.bias=tab.bias[order(tab.bias$msrun_ID, tab.bias$rt),]
	tab.bias=tab.bias[!is.na(tab.bias$rt),] 
	tab.bias$correc=NA
	compteur=unique(tab.bias$msrun_ID)

	pdf("intensity_bias.pdf")
		for (i in 1:length(compteur)) {
		# compute the correction factor
			select.samp= tab.bias$msrun_ID==compteur[i]	
			select.na=is.na(tab.bias$bias)
				
			cour=tab.bias[select.samp,]
			cour.nomiss=tab.bias[select.samp & !select.na,]

			rt.min=min(cour.nomiss$rt)
			rt.max=max(cour.nomiss$rt)
			bias.min=min(cour.nomiss$bias)
			bias.max=max(cour.nomiss$bias)
			plot(cour.nomiss$rt,cour.nomiss$bias,main=paste("msrun n°", compteur[i], sep=""),xlab="RT",ylab="Bias (log)",xlim=c(rt.min,rt.max),ylim=c(bias.min,bias.max))
			yyy=smooth.spline(cour.nomiss$rt,cour.nomiss$bias,spar=0.5)
			lines(yyy$x,yyy$y,col="red",type="l")
			indice=match(tab.bias$rt[select.samp & !select.na], yyy$x)
			tab.bias$correc[select.samp & !select.na] = yyy$y[indice]	

		# propagation to the peptiz that were missing in the reference msrun
			smo=tab.bias$correc[select.samp] 
			j=1
			while(is.na(smo[j])) {smo[j]=0; j=j+1}
			while (j < length(smo)) {
				while(!is.na(smo)[j] & j<length(smo)) {last=smo[j]; j=j+1}
				while(is.na(smo[j]) & j<length(smo)) {smo[j]=last; j=j+1}
			}  
			smo[length(smo)]=smo[length(smo)-1] 
			tab.bias$correc[select.samp] =smo
			print(i)
		}
	dev.off()

	### merging tab.quanti and tab.bias
	indice=match(paste(tab.quanti$msrun_ID, tab.quanti$peptiz), paste(tab.bias$msrun_ID, tab.bias$peptiz))
	tab.quanti$correc=tab.bias$correc[indice]
	tab.quanti$logQnorm=tab.quanti$logQ+tab.quanti$correc


############# Post-normalization filters on peptiz

	### filter peptides identified as a mutated version of the peptide sequences present in the interrogated database
	mut.motifs=strsplit(as.character(tab.mut$Modifs), "@", fixed=TRUE)
	long=unlist(lapply(mut.motifs, length))
	select=long>1 #mutated peptides are those with "pm@" in the "Modifs" column 
	SAP=tab.mut[select,] 
	pepSAP=unique(SAP$Peptide)
	pepz.mut=unique(tab.seq$peptiz[tab.seq$peptide %in% unique(SAP$Peptide)])
	tab.quanti=tab.quanti[!tab.quanti$peptiz %in% pepz.mut,] #17,099,482 obs.

	### filter shared peptides
	count = as.data.frame(table(tab.prot$peptide))
	common.peptide=count$Var1[count$Freq>1] 
	tab.seq=tab.seq[tab.seq$msrun_ID %in% unique(tab.quanti$msrun_ID),]
	common.peptiz=unique(tab.seq$peptiz[tab.seq$peptide %in% common.peptide]) 
	tab.quanti=tab.quanti[!tab.quanti$peptiz %in% common.peptiz,] #12,927,341 obs.

	### add the protein information in tab.quanti
	tab.temp=tab.seq[tab.seq$peptiz %in% unique(tab.quanti$peptiz),] 
	tab.temp=merge(tab.temp, tab.prot, "peptide")
	tab.quanti=merge(tab.quanti, unique(tab.temp[,c("peptiz", "protein")]), "peptiz") #12,927,341 obs., 19,009 peptiz 

	### filter peptiz showing too many missing inensity values
	count=table(tab.quanti$peptiz)
	nb.sample=length(unique(tab.quanti$sample_ID))
	names.pep.rep=names(count[count>(nb.sample-nb.sample*0.1)]) #7,554 peptiz
	tab.quanti=tab.quanti[tab.quanti$peptiz %in% names.pep.rep,] #7,161,539 obs, 
	
	### filter peptiz poorly correlated to the other peptiz belonging to the same protein
	# we want a protein to be quantified by at least two different peptides
	temp=unique(tab.quanti[,c("protein", "peptiz")])
	spli=strsplit(temp$peptiz, "-")
	peptide=NULL
	for (i in 1:nrow(temp)){
		peptide=c(peptide, spli[[i]][1])

	}
	temp$peptide=peptide 
	temp=unique(temp[,c("peptide", "protein")])
	count=table(temp$protein)
	prot.rep=names(count[count>1]) #1260 prot
	tab.quanti=tab.quanti[tab.quanti$protein %in% prot.rep,] #6,490,754 obs.

	# compute correlations between peptides 
	tab.quanti$genocond=paste(tab.quanti$hybrid_ID, tab.quanti$watering_condition, sep="-")
	proteines = unique(tab.quanti$protein)
	test.corr=NULL
	for (i in 1:length(proteines)){
 		sub=tab.quanti[tab.quanti$protein==proteines[i],]
 		mytab=tapply(sub$logQnorm,list(sub$genocond,sub$peptiz),FUN=mean,na.rm=T) # we consider intensities averaged across replicates
 		for (j in 1:length(unique(sub$peptiz))){
			for (k in 1:length(unique(sub$peptiz))){
				if (nrow(na.omit(cbind.data.frame(mytab[,j], mytab[,k])))>6){ # we want at least 6 points to compute a correlation
					temp=cor.test(mytab[,j], mytab[,k])
					test.corr=rbind.data.frame(test.corr, cbind.data.frame(peptiz1=colnames(mytab)[j], peptiz2=colnames(mytab)[k], protein=proteines[i], pval=temp$p.value, r=temp$estimate))
				}
			}
		}
	print(i)
	}

	fdr=p.adjust(test.corr$pval, method="BH")
	test.corr=cbind.data.frame(test.corr, fdr=fdr)# 73867 lignes

	# find the peptide that will be used as the best representative of the protein 
	temp=test.corr[test.corr$fdr<0.01 & test.corr$r!=1,] 
	temp=drop.levels(temp)
	count=as.data.frame(table(temp$protein, temp$peptiz1))
	count=count[!count$Freq==0,]
	prot=unique(count$Var1) #1217 prot	
	thebest.pep=NULL
	for ( i in 1:length(prot)){
		sub.count=count[count$Var1==prot[i],]
		max.count=sub.count[sub.count$Freq==max(sub.count$Freq),]
		thebest.pep=c(thebest.pep, as.character(max.count[1,2])) 
		print(i)
	}
	tab.best.pep=test.corr[test.corr$peptiz1 %in% thebest.pep,]
	tab.best.pep=drop.levels(tab.best.pep) #6751 obs et pepz2, 1217 prot et pepz1


	# compute the correlations to the reference peptide
	signif=tab.best.pep[tab.best.pep$fdr<0.01 & tab.best.pep$r>0.3,] 
	signif=drop.levels(signif)#4461 obs, 1217 prot et pepz1, 4461 pepz2
	tab.quanti=tab.quanti[tab.quanti$peptiz %in% unique(signif$peptiz2),] #4249269, 4461 pepz, 1217 prot
	

	###  post-peptide filter: remove the proteins quantified by less than two different peptides
	temp=unique(tab.quanti[,c("protein", "peptiz")])
	spli=strsplit(temp$peptiz, "-")
	peptide=NULL
	for (i in 1:nrow(temp)){
		peptide=c(peptide, spli[[i]][1])

	}
	temp$peptide=peptide #4196 pep, 4461 pepz, 1217 prot
	temp=unique(temp[,c("peptide", "protein")])
	count=table(temp$protein)
	prot.rep=names(count[count>1]) #974 prot
	tab.quanti=tab.quanti[tab.quanti$protein %in% prot.rep,] #4011567 obs, 974 prot, 4208 peptiz
