## supplemental methods for "A systems genetics approach reveals environment-dependent associations between SNPs, protein co-expression and drought-related traits in maize"

### Plant material and experiment

A diversity panel of maize hybrids obtained by crossing a common flint parent (UH007, paternal parent) with 254 dent lines was used. The experiment was carried out in the phenotyping platform PhenoArch (Cabrera-Bosquet et al. 2016) ((https://www6.montpellier.inra.fr/lepse_eng/M3P/PHENOARCH-platform) hosted at the Montpellier Plant Phenotyping Platforms (https://www6.montpellier.inra.fr/lepse_eng/M3P). Plants were sown on May 14, 2012 and grown in 9L pots (0.19 m diameter, 0.4 m high) filled with a 30:70 (v/v) mixture of a clay and organic compost. Three seeds per pot were sown at 0.025 m depth and thinned to one per pot when leaf three emerged. Two levels of soil water content were imposed, namely retention capacity (well-watered, soil water potential of −0.05 MPa) and water deficit (soil water potential from −0.45 MPa). The weight of water in each pot was calculated at the beginning of the experiment from the weight of soil and measured soil water content. Soil water content in pots was maintained at target values by watering each pot three times per day, using watering stations made up of weighting terminals with 1 g accuracy (ST-Ex, Bizerba, Balingen, Germany) and high-precision pump-watering stations (520 U, Watson Marlow, Wilmington, MA, USA). The greenhouse temperature was maintained at 26 ± 3 °C during the day and 18 ± 1 °C during the night. Supplemental light was provided either during daytime when external solar radiation was below 300 W $m^{-2}$ or to extend the photoperiod by using 400 W HPS Plantastar lamps (OSRAM, Munich, Germany) with 0.4 lamps $m^{-2}$. Each hybrid was replicated 4 and two times for the well-watered and water deficit treatments, respectively.

Sampling was performed at the pre-flowering stage (between June 19 and June 22, 2012) in two replicates per hybrid and water condition. For each sampled plant, 35 to 45 mg of fresh material was taken in a 2 mL tube containing two iron metal beads (5 mm diameter) by punching ten patches

(5 mm diameter) in the mature area of the last ligulated leaf. The tubes were frozen in liquid nitrogen immediately after sampling and stored at -80°C until protein extraction.

**Protein extraction and digestion**

Sampled leaf patches were turn into powder by shaking frozen sample tubes twice at 20 Hz for 20 seconds using TissueLyzer II (Quiagen, Courtaboeuf, France). Beads were removed using a magnet. Proteins were precipitated by incubating the leaf powder in 1.2 ml of an ice-cold solution of acetone containing 10% of trichloroactetic acid and 0.07% β-mercaptoethanol for 1h20 at -20°C. After centrifugation (10 min, 0°C, 14 000 rpm), the supernatants were removed and the protein extracts were washed by incubation in 1.2 ml of 0.07% β-mercaptoethanol in acetone (1h, -20°C). This step was repeated twice. After the last washing, proteins were dried in a vacuum centrifuge and stored at -80°C until solubilization.

Dried protein pellets were solubilized in 100 µl of a solution containing 6 M of urea, 2 M of thiurea, 10 mM of dithiothreitol (DTT), 30 mM of TrisHCl pH 8.8 and 0.1% of Zwitterionic Acid Labile Surfactant I (ZALS I, Protea Bioscience, Morgantown, USA). Protein powders were mixed in the buffer using a metal spatula before vortexing the tubes for 3 min. Remaining cellular debris were segregated from soluble proteins by centrifugation (12,500 rpm, 25 min, room temperature). Protein concentrations were determined using the PlusOne 2-D Quant kit (GE Healthcare, Little Chalfont, UK) and adjusted to 4 $\mu g.\mu l^{-1}$ prior to digestion.

Digestion was performed in 0.2 ml strip tubes from 10 µl of diluted proteins. Proteins were incubated one hour at room temperature for reduction by the 10 mM DTT present in the buffer. Thereafter, proteins were alkylated one hour in 40 mM iodoacetamide (room temperature in the dark) and diluted with 50 mM ammonium bicarbonate to decrease total urea and thiourea concentration to 0.77 M. Overnight digestion was performed at 37°C with 1/50 (w/w) trypsin (Promega, Charbonnières-les-Bains, France) and stopped by acidification (1% total volume of trifluoroacetic acid, TFA). The resulting peptides were desalted on solid phase extraction using polymeric C18 columns (strata XL 100 µm, ref 8E-S043-TGB; Phenomenex, Le Pecq, France) as follows. Peptides were first diluted in 3% ACN and 0.06% acetic acid in water (washing buffer) up to a final volume of 500 µl. Then, they were loaded onto cartridges previously conditioned with 500 µl of ACN and rinsed three times with 500 µl of washing buffer. Peptides were rinsed three times with 500 µl of washing buffer and eluted twice by adding 300 µl of 40% ACN and 0.06% acetic acid. To finish, eluted peptides were speed-vac dried and suspended in a solution containing 2% ACN, 0.05% TFA and 0.05 % formic acid.

**Parameters of data-dependent acquisition**

For mass-spectrometry analysis, the following data-dependent acquisition steps were performed: (1) MS scan (mass-to-charge ratio (m/z) 400 to 1400, 70,000 resolution, profile mode), (2) MS/MS (isolation window of 3 m/z, 17,500 resolution, collision energy = 27%, profile mode). Step 2 was repeated for the eight major ions detected in step 1 with a charge of 2 or 3. Dynamic exclusion was set to 40 s. Xcalibur raw datafiles were transformed to mzXML open source format using msconvert software in the ProteoWizard 3.0.3706 package (Kessner et al. 2008). During conversion, MS and MS/MS data were centroided.

**Parameters of peptide identification and protein inference**

For peptide identification, enzymatic cleavage was declared as a trypsin digestion with one possible misscleavage. Cystein carboxyamidomethylation and methionine oxidation were set to static and possible modifications, respectively. Precursor mass error was set to 10 ppm and fragment mass tolerance was set to 0.02 Da. In refine mode, a second search was performed with the same settings, except that protein N-ter acetylation was added as a potential modification and that the point mutations option was activated to detect possible single amino acid changes in the peptide sequences. Only peptides with an E-value smaller than 0.01 were reported.

For protein inference, only the proteins identified with a minimum of two peptides were considered as valid. Protein inference was performed using all samples together.

The false discovery rate (FDR) was assessed from searches against a decoy database (using the reversed amino acid sequence for each protein).

**Parameters of peptide quantification**

Peptide quantification was performed using MassChroQ version 2.1.0 (Valot et al. 2011) based on extracted ion chromatograms (XIC) with the following parameters: "ms2_1" alignment method, tendency_halfwindow of 10, MS1 smoothing halfwindow of 0, MS2 smoothing halfwindow of 15, "quant1" quantification method, XIC extraction based on max, min and max ppm range of 10, anti-spike half of 5, background half median of 5, background half min max of 20, detection thresholds on min and max at 30 000 and 50 000, respectively, peak post-matching mode, "ni min abundance" of 0.1.

**Filtering and normalizing peptide intensity data**

R script for filtering and normalizing peptide intensity data can be found in Supplemental Material. We first removed the peptides ions showing standard deviations of retention time >15 s, which may arise from mis-identifications. Intensity normalization was subsequently performed to take into account possible global quantitative variations between LC-MS/MS runs. For this, we used a local normalization method described in Millan-Oropeza et al. (2017) and adapted from (Lyutvinskiy et al. 2013). In brief, intensity deviation between a given sample and a sample chosen as reference was computed for each peptide charge quantified in both samples. The values of intensity deviation thus obtained were ordered according to the peptides' retention time and smoothed using the smooth.spline function in R. The smoothed values of intensity deviation were used as correction factors to normalize the sample. The intensities of the peptides charges that were present in the sample to be normalized and absent from the reference sample were corrected considering that intensity deviation was similar for temporally neighboring peptides.

We then removed the peptides shared between several proteins as well as the peptides for which both the unmodified form and a mass modification corresponding to an amino acid change was detected by the point mutation option of X!Tandem. These mutated peptide forms represent allelic versions of the sequences present in the searched protein database. In heterozygous genotypes, each allelic version produces its own MS signal, so that their measured intensities are not representative of the total protein abundance. We also removed the peptide ions presenting more than 10% missing values and those showing inconsistent intensity profiles. To this end, we computed Pearson correlations between log-intensities averaged across replicates for each pair of peptide ions belonging to the same protein. The peptide ion with the highest number of significant correlations (*P-value* < 0.01 after adjustment for multiple testing (Benjamini and Hochberg 1995) was chosen as a reference for the protein. The peptide ions showing non-significant correlation to the reference (adjusted *P-value* >= 0.01) or whose coefficients of correlation to the reference were inferior to 0.3 were removed.

**XIC-based protein quantification and differential analysis**

For each protein, we modeled the peptide data using the following mixed-effects model derived from Blein-Nicolas et al. (2012):

$$I'_{ijkl} = \mu + G_j + E_k + (GxE)_{jk} + R_{l(k)} + P_i + \theta_{jkl} + \varepsilon_{ijkl} \quad (1)$$

where $I'_{ijkl}$ is the corrected, normalized log-intensity measured for peptide *i* in genotype *j*, watering condition *k* and replicate *l;*

$\mu$ is the mean intensity for a given protein;

$G_j$ is the effect of the genotype *j;*

$E_k$ is the effect of the watering condition *k;*

$(GxE)_{jk}$ is the effect of the genotype *j* x watering condition *k* interaction;

$R_{l(k)}$ is the effect of the replicate *l* nested in the watering condition *k;*

$P_i$ is the effect of the peptide *i;*

$\Theta_{jkl} \sim \mathcal{N}(0, \sigma_\Theta^2)$ is the random technical variation due to handling and injection in the mass spectrometer of the sample *jkl*;

$\varepsilon_{ijkl} \sim \mathcal{N}(0, \sigma_\varepsilon^2)$ is the residual error.

Model parameters were estimated by maximizing the restricted log-likelihood (REML method) and the differential protein abundance analysis was performed by analysis of variance (ANOVA). The resulting *P-value*s were adjusted for multiple testing by the Benjamini-Hochberg procedure (Benjamini and Hochberg 1995). Proteins showing significant abundance variation were detected based on the following criteria: adjusted *P-value* < 0.05, WD/WW ratio or maximum ratio between genotypes > 1.5 or < 0.66.

To subsequently perform GWAS at the protein level, we estimated protein abundances in each watering condition using the following model:

$$I'_{ijkl} = \mu_k + G_{jk} + R_{lk} + P_{ik} + \theta_{jkl} + \varepsilon_{ijkl} \quad (2)$$

where $\mu_k$ is the mean intensity obtained for a given protein in the watering condition *k*;

$G_{jk}$ is the effect of the genotype *j* in the watering condition *k;*

$R_{lk}$ is the effect of the replicate *l* in the watering condition *k;*

$P_{ik}$ is the effect of the peptide *i* in the watering condition *k.*

Protein abundances were computed as adjusted means as follows:

$$A_{jk} = \mu_k + G_{jk} \quad (3)$$

**Spectral counting-based protein quantification and differential analysis**

Normalization of spectral counts was performed as follows:

$$Anorm_{ps} = \frac{A_{ps}}{\sum_{n=1}^{P} A_{ns}} x \frac{\sum_{m=1}^{S} \sum_{n=1}^{P} A_{nm}}{S} \quad (4)$$

where $A_{ps}$ is the abundance of protein $p$ in sample $s$;

$P$ is the number of quantified proteins;

$S$ is the number of samples.

As for XIC-based quantification, we corrected the batch effect by fitting a linear model to square-root transformed, normalized protein abundances. Differential protein abundance analysis was subsequently performed by analysis of variance (ANOVA) using the following mixed effect model:

$$A'_{jkl} = \mu + G_j + E_k + (GxE)_{jk} + \alpha_l + \epsilon_{jkl} \quad (5)$$

where $A'_{jkl}$ is the corrected, normalized, square-root transformed abundance obtained for a given protein in genotype $j$, watering condition $k$ and replicate $l$;

$\mu$ is the mean abundance for the protein;

$G_j$ is the effect of the genotype $j$;

$E_k$ is the effect of the watering condition $k$;

$(GxE)_{jk}$ is the effect of the genotype $j$ x watering condition $k$ interaction;

$\alpha_l \sim \mathcal{N}(0, \sigma_{\alpha}^2)$ is the random effect of the replicate $l$;

$\varepsilon_{ijkl} \sim \mathcal{N}(0, \sigma_{\varepsilon}^2)$ is the residual error.

Estimation of the model parameters, differential analysis and *P-value* adjustment were performed as described above. Finally, for GWAS, we estimated the protein abundances separately in each watering condition with a mixed model derived from (5) and including only a fixed effect of the genotype and a random effect of the replicate. Protein abundances were computed as adjusted means as in (3).

**Computing of broad sense heritability**

For each protein, the broad sense heritability of abundance was computed in each of the two watering conditions from a mixed effects model as follows. For the proteins quantified by the XIC-based approach, abundances were estimated in each sample as adjusted means from (1), by excluding the peptide effect $P_{ik}$ and the random sample effect $\Theta_{jkl}$. For the proteins quantified by the

SC-based approach, the corrected, normalized, square-root transformed abundances were used. Protein abundances were then modeled as follows:

$$A_{jkl} = \mu_k + \beta_{jk} + \gamma_{kl} + \epsilon_{jkl} \qquad (6)$$

where $A_{ijk}$ is the abundance estimated for a given protein in the genotype *j*, the replicate *i* and the watering condition *k*

$\beta_{jk} \sim \mathcal{N}(0, \sigma_{\beta}^2)$ is the random effect of the genotype *j* in the watering condition *k*

$\gamma_{kl} \sim \mathcal{N}(0, \sigma_{\gamma}^2)$ is the random effect of the replicate *l* in the watering condition *k.*

Heritability was subsequently computed as

$$H^2 = \frac{\sigma_\beta^2}{\left(\sigma_\beta^2 + \sigma_\gamma^2 / N\right)} \qquad (7)$$

where *N* is the number of replicates.

**Parameters of WGCNA analysis**

Protein co-expression analysis was performed using the WGCNA R package (Langfelder and Horvath 2008) with the following parameters. The softpower parameter was set at 2. Adjacency and topological ovelap matrices were both unsigned. Protein modules were constituted with a minimum module size set at 20 and a control over sensitivity splitting set at 4. The minimum height for merging modules was set at 0.25. The other parameters were left at default values. For each module, eigengene was computed as described in the WGCNA R package, *i.e.* as the first principal component of the matrix of protein abundances of the corresponding module.
